# Decorin-mediated Extracellular Matrix Stabilization Maintains Cartilage Integrity During Aging

**DOI:** 10.64898/2026.07.28.741218

**Authors:** Mingyue Fan, Thomas Li, Aanya Mohan, Biao Han, Michael Newton, Bryan Kwok, Yuchen Liu, Sung Yeon Kim, Chao Wang, Jiaqi Xiang, Ling Qin, Renato V. Iozzo, David E. Birk, X. Lucas Lu, Claudia Loebel, Tristan Maerz, Robert L. Mauck, Lin Han

## Abstract

Aging is a primary risk factor for osteoarthritis (OA), yet the mechanisms that preserve extracellular matrix integrity in long-lived connective tissues remain poorly defined. The function of articular cartilage depends on maintaining extracellular matrix integrity over a lifetime of mechanical use. Here we identify failure of matrix stabilization as an initiating mechanism of age-associated OA. Cartilage-specific deletion of decorin in mature mice disrupted the superficial collagen II fibrillar network, reduced aggrecan retention, and impaired poroelastic fluid pressurization by 9 months of age, preceding substantial transcriptional changes in resident chondrocytes. With advancing age, these matrix defects culminated in cartilage erosion, fibrotic remodeling, and spontaneous OA by 18 months. Mechanistically, decorin attenuated force-induced collagen II fibril realignment and reinforced the superficial fibrillar network, preserving matrix architecture under sustained physiological loading. These findings establish decorin-mediated extracellular matrix stabilization as a critical determinant of cartilage longevity and support a matrix-first model of age-associated degeneration.

## INTRODUCTION

Aging is a primary risk factor for osteoarthritis (OA), the most prevalent musculoskeletal disorder affecting 595 million people (7.6% of the global population worldwide in 2023).^1^ Approximately one in three adults over age 65 develops symptomatic OA.^2^ Beyond its clinical burden, OA represents a prototypical example of age-associated load-bearing tissue degeneration, characterized by progressive and irreversible breakdown of articular cartilage.^3^ Articular cartilage is a specialized avascular connective tissue that covers diarthrodial joint surfaces and enables joint motion, efficient load transmission and shock absorption.^4^ The limited intrinsic regenerative capacity of cartilage renders it particularly vulnerable to cumulative age-related damage.^5^ Effective clinical management of aging-associated OA is currently unavailable, in part due to the low regenerative capacity of cartilage and an incomplete understanding of the molecular mechanisms driving cartilage aging.^6^

Cartilage aging reflects coordinated changes in both chondrocyte cellular function and extracellular matrix (ECM) integrity, and disruptions in their dynamic reciprocal interactions.^7^ Extensive efforts have focused on elucidating aging-associated cellular mechanisms of tissue decline, including DNA damage and senescence,^8–10^ telomere attrition,^11^ mitochondrial dysfunction,^12^ impaired antioxidant defense,^13^ alterations in sex hormones,^14^ reduced autophagy with increased apoptosis,^15^ chronic inflammation,^16^ as well as dysregulated anabolic-catabolic balance.^17,18^ These insights have motivated regenerative strategies aimed at targeting these molecular processes towards cellular rejuvenation.^19^ However, articular cartilage represents a matrix-dominant tissue in which long-term function depends less of cellular turnover and more on the sustained preservation of extracellular structural architecture throughout lifespan.^20^ Some studies even indicate that a loss of cells may be protective to cartilage damage following injury through blocking aggravated catabolic cellular factors.^21^ In tissues with low cellularity and limited regenerative renewal such as cartilage, failure of matrix-stabilizing mechanisms may constitute an initiating driver of age-associated degeneration. Whether progressive destabilization of ECM organization can precede and independently trigger cartilage aging remains poorly defined.^6^ More broadly, establishing how extracellular proteostasis is actively maintained across the lifespan is essential for understanding the origins of matrix-associated diseases^22^ including OA.

Cartilage ECM is a hydrated composite mainly consisting of a collagen II fibrillar network entrapping the large proteoglycan aggrecan.^23^ This architecture confers the biomechanical function necessary for withstanding physiological joint loading while at the same time providing a finely tuned translation of these physical forces to regulate mechanosensitive signaling that sustains chondrocyte homeostasis.^23,24^ During aging, these two primary ECM constituents undergo progressive remodeling at multiple scales. Tissue-level changes include reduced cartilage thickness, decreased sulfated glycosaminoglycan (sGAG) content, decreased tensile^25^ and shear^26^ moduli (in human tissue), as well as lower indentation moduli (in murine cartilage).^27^ At the molecular level, aggrecan exhibits reduced GAG side chain length, density, altered sulfation patterns,^28^ and increased fragmentation, leading to a lower fixed charge density and compressive modulus.^29^ Collagen II fibrils display increased degradation^30^ and accumulation of non-enzymatic advanced glycation end-product (AGE)-mediated cross-links,^31^ which are characteristic of aging ECMs across tissues. Such changes were hypothesized to increase fibril stiffness and brittleness, and have been proposed to perturb chondrocyte mechanotransduction, potentially amplifying aging-associated functional decline.^32–34^ Although collagen II and aggrecan constitute the principal structural framework of cartilage, their organization and long-term stability depend on coordinated interactions with other regulatory matrix molecules.^35^ The identification of ECM regulators that preserve cartilage structural integrity during aging, and the mechanisms by which they modulate age-dependent matrix remodeling, remain largely undefined. This represents a critical gap in our understanding of how extracellular proteostasis and matrix architecture are maintained in long-lived connective tissues.^36^

Decorin, a small leucine-rich proteoglycan, is a candidate regulator of matrix homeostasis in aging cartilage. Although classically recognized for modulating collagen I fibrillogenesis,^37^ decorin is present in human cartilage at a molar concentration (≈ 15 nmol/mL) comparable to that of aggrecan (≈ 20 nmol/mL).^38^ This molar ratio suggests that decorin may play a substantial structural role within the cartilage matrix. Indeed, our prior studies have established that decorin preserves aggrecan content during postnatal maturation^39,40^ and mitigates matrix degeneration following joint injury,^41,42^ supporting its contribution to tissue-level biomechanics^39^ and the pericellular matrix (PCM) micromechanics that regulate chondrocyte mechanosensing.^43^ However, whether decorin actively maintains cartilage matrix integrity during aging, and whether its loss initiates progressive matrix destabilization, remains unknown. Given its sustained expression throughout the human lifespan,^44–46^ we hypothesize that decorin functions as a long-term regulator of extracellular proteostasis that is required for the structural resilience that characteristics cartilage ECM. Here, using a cartilage-specific decorin knockout murine model, we assessed how post-maturation ablation of decorin influences cartilage aging across structural, biomechanical, cellular and molecular scales. We show that decorin deficiency triggers early remodeling of cartilage surface fibrillar network and loss of its matrix energy-dissipative function, preceding overt osteoarthritic structural degeneration and major transcriptional changes in chondrocytes. This progressive matrix destabilization ultimately culminates in cartilage erosion, aberrant fibrotic remodeling, disrupted cell signaling, and infiltration of non-resident cell populations. At the cellular and molecular levels, we further show that decorin attenuates force-induced collagen II fibril realignment and reinforces the superficial collagen II fibrillar architecture. Together, these findings reveal a matrix-centered mechanism governing age-associated cartilage degeneration, and position decorin-mediated ECM stability as a primary determinant of cartilage longevity.

## RESULTS

### Loss of Decorin Induces Early Cartilage Surface Remodeling and Accelerates Age-associated Cartilage Erosion

In human cartilage from aged donors, we showed that decorin was present throughout tissue depth in both normal and early OA specimens, with a preferential localization in the pericellular domain (Fig. 1a), consistent with the literature.^44–46^ Given this persistent and spatially defined distribution, we first tested whether decorin is required for cartilage ECM maintenance during aging. We generated cartilage-specific inducible *Dcn* knockout mice (*Dcn^f/f^/AcanCre^ER^*, or *Dcn^cKO^*), induced *Dcn* gene ablation at 3 months of age, and analyzed the joint phenotype at 9 and 18 months, following confirmation of efficient decorin deletion in cartilage (Fig. 1b). These ages correspond to young adult (20-30 years), middle aged (35-45 years) and old (56-69 years) humans, respectively. At 9 months, *Dcn^cKO^* cartilage exhibited a moderate reduction in sGAG staining (Fig. 1c), consistent with our prior findings that decorin stabilizes aggrecan during postnatal maturation.^39^ Despite this reduction, cartilage thickness, cellularity and Mankin scores were comparable to controls (Fig. 1d-f), indicating that early, moderate loss of aggrecan has not yet progressed to overt OA pathology. However, notably, *Dcn^cKO^* cartilage developed a distinct sGAG-depleted layer on the femoral cartilage surface (Fig. 1g, arrowhead). This region displayed pronounced birefringence under polarized light microscopy (PLM, Fig. 1h), indicative of collagen fibril thickening and alignment. Scanning electron microscopy (SEM) confirmed that the condylar surface became dominated by densely packed, highly aligned collagen fibrils along the mediolateral split-line axis (Fig. 1i,j). Quantitative analysis revealed significantly increased fibril alignment (von Mises concentration *κ*, Fig. 1k) and fibril diameter compared to controls (Fig. 1l). In contrast, the control cartilage surface retained the transversely isotropic random meshwork characteristic of normal hyaline cartilage^47^ (Fig. 1i-l). Notably, no surface or tissue phenotypes were detectable in 4-month-old *Dcn^cKO^* joints following the induced *Dcn* ablation at 3 months (Extended Data Fig. 1a-d), suggesting that the phenotype at 9 months reflects progressive, cumulative effects of decorin loss during post-maturation cartilage maintenance. Together, these findings suggest that loss of decorin induces extracellular structural remodeling that emerges during early aging in the absence of overt degenerative pathology.

**Figure 1.**
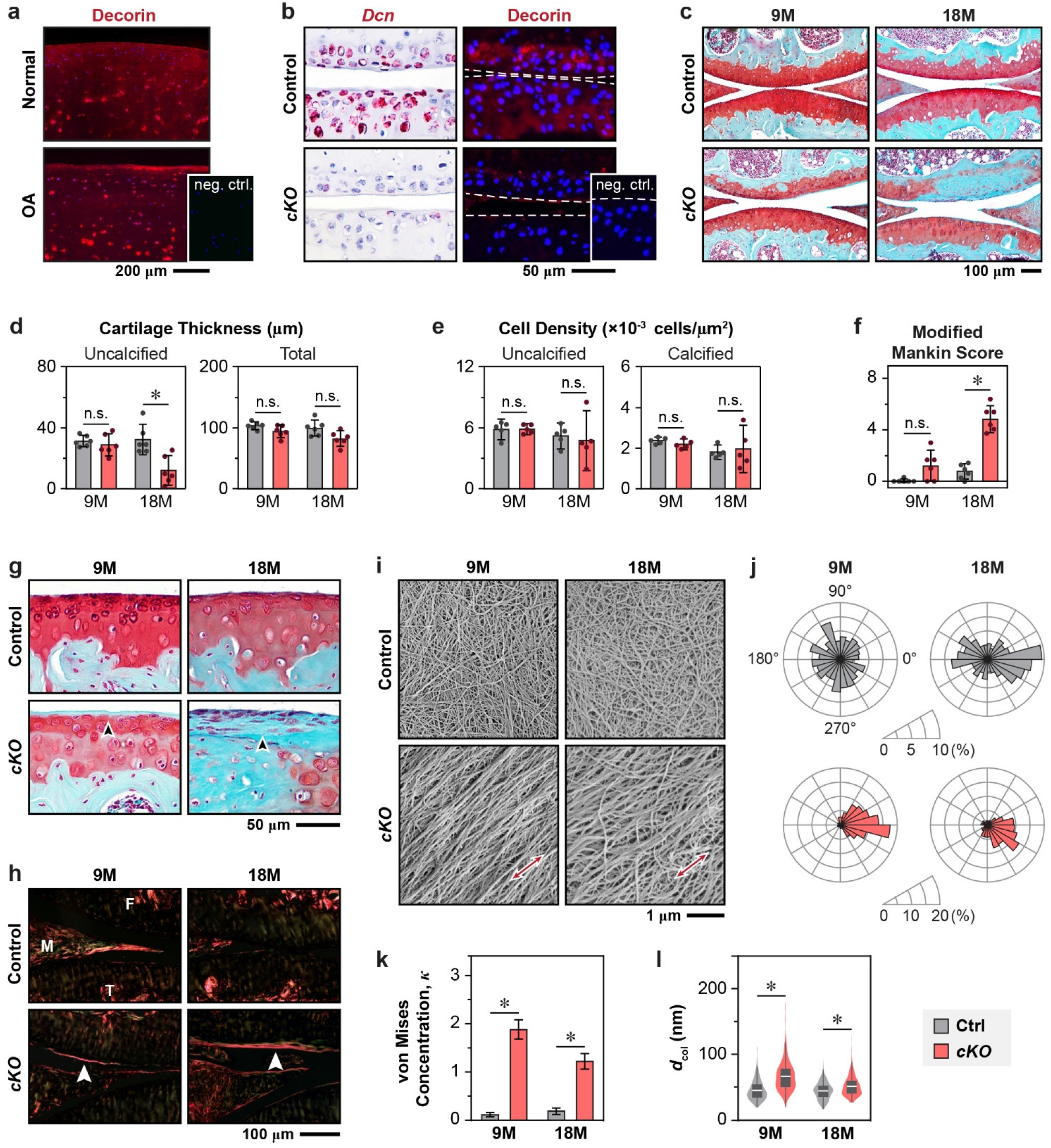
Cartilage-specific ablation of decorin results in early cartilage surface fibril remodeling and accelerates age-associated cartilage erosion. **a)** Decorin immunofluorescence (IF) images illustrate the presence of decorin in articular cartilage from both normal (female, 61 years of age) and osteoarthritis (OA, female 55 years of age) human donors. **b)** RNAscope *in situ* hybridization for *Dcn* mRNA and decorin IF validate decorin ablation in 9-month-old (9M) cartilage-specific inducible *Dcn* knockout mice (*Dcn^flox/flox^/AcanCre^ER^*, or *cKO*) mice compared to age-matched controls, following the induced ablation at 3 months of age (red: decorin; blue: DAPI, *n* = 5). **c)** Representative Safranin-O/Fast Green staining of sagittal knee joint sections at the primary mechanical loading region shows progressive structural disruption and sulfated glycosaminoglycan (sGAG) loss in 9M and 18M *cKO* cartilage compared to controls (*n* ≥ 8). **d–f)** Quantification of d) cartilage thickness, e) chondrocyte cell density and f) modified Mankin scores from femoral condyle histological sections (mean ± 95% CI; *: *p* < 0.05; n.s.: not significant; *n* ≥ 5. Each data point represents the average value measured from one animal). **g)** High-magnification images of femoral condyle cartilage highlight early surface remodeling at 9M and full erosion of uncalcified cartilage at 18M in *cKO* joints (black arrowheads). **h)** Polarized light microscopy (PLM) images of Picrosirius Red (PSR)-stained sections illustrate increased collagen fiber alignment on *cKO* cartilage surface (*n* = 4). **i)** Scanning electron microscopy (SEM) images show the formation of aligned collagen fibrils along the mediolateral split-line direction (red arrows) on *cKO* cartilage surface (*n* ≥ 5). **j–l)** Quantitative analysis of collagen fibril organization from SEM images, including j) rose plots of fibril orientation distribution, where 0° corresponds to the mediolateral split-line direction, k) degree of fibril alignment as denoted by the von Mises concentration, *κ* (mean ± 95% CI), and l) box-and-whisker plot of collagen fibril diameter, *d*_col_, distribution (*n* = 400 fibrils pooled from *n* = 4 animals per group). A complete list of quantitative and statistical outcomes is summarized in Supplementary Tables S1 and S2.

In contrast to the early time points, by 18 months, *Dcn^cKO^*joints exhibited pronounced degenerative changes (Fig. 1c). The femoral condyle showed near-complete erosion of uncalcified cartilage (Fig. 1c, d), accompanied by significantly higher Mankin scores (Fig. 1f). This eroded surface was covered by an abnormal pannus-like fibrotic layer lacking sGAG staining (Fig. 1c,g), which was populated by morphologically aberrant cells that were distinct from resident chondrocytes (Fig. 1g), and enriched in thick, aligned collagen fibers (Fig. 1h-l). These phenotypic changes were observed in both male and female mice (Extended Data Fig. 1e-f), suggesting a sex-independent role of decorin in cartilage aging.

The cartilage-specific knockout model did not exhibit major alterations in other joint tissues. We found comparable subchondral bone plate thickness and trabecular bone structure between genotypes at both ages, although increased ossification of meniscal horns, which characterizes advanced OA in mice^41^ was observed in 18-month-old *Dcn^cKO^*mice (Extended Data Fig. 2). Also, the synovium remained appreciable decorin expression for both genotypes. Synovitis scores were largely uncharged, except for mild hyperplasia at 18 months (Extended Data Fig. 3). These minimal-to-mild changes observed in other tissues indicate that the observed cartilage phenotypes represent the primary role of decorin in cartilage maintenance, instead of secondary effects arising from osteal or synovial tissues.

### Matrix Destabilization Impairs Poroelastic Function Prior to Overt Structural Degeneration

We next queried how matrix remodeling resulting from a loss of decorin impaired the biomechanical function of cartilage. Applying atomic force microscopy (AFM)-nanoindentation, we first quantified the effective indentation modulus, *E*_ind_, a measure of tissue load-bearing capacity. At 9 months of age, *E*_ind_ did not differ significantly between *Dcn^cKO^*and control cartilage (Fig. 2a). This finding suggests that the reduction in sGAG content may be mechanically offset by the surface stiffening effect from increased collagen fibril alignment, resulting in no net change in quasi-static modulus at this stage. In contrast, by 18 months, *Dcn^cKO^* cartilage exhibited substantial stiffening relative to controls (Fig. 2a), consistent with the deposition of a dense, fibrotic surface layer enriched in aligned collagen fibers (Fig. 1g-i). Thus, while early decorin loss did not measurably alter tissue modulus, progressive matrix remodeling ultimately resulted in abnormal surface stiffening.

**Figure 2.**
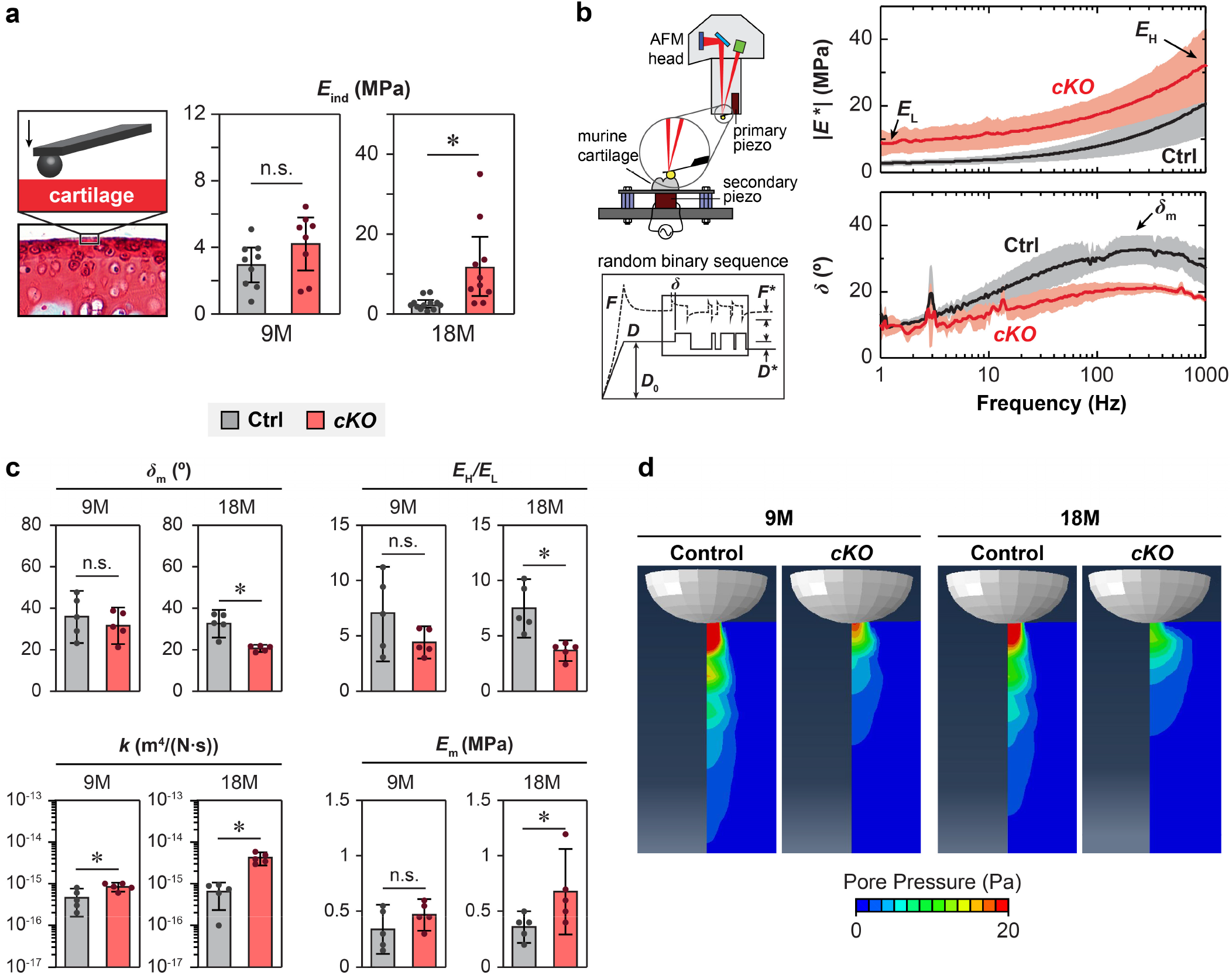
Cartilage-specific ablation of decorin leads to impaired elastic and poroelastic mechanical properties of cartilage. **a)** Left panel, schematic of AFM-nanoindentation on the surface of femoral condyle cartilage. Right panel, comparison of the indentation modulus (*E*_ind_) between control and *cKO* cartilage at 9 and 18 months (9M and 18M) (*n* ≥ 8). **b)** Left panel, schematic of the custom-built nanorheometer and representative force-displacement profiles incorporating nanoscale dynamic oscillations in the form of a random binary sequence (RBS) superimposed onto a static 60-second ramp-and-hold period. Right panel, representative frequency spectra of dynamic modulus (|*E*\*|) and phase angle (*δ*) from control and *cKO* cartilage at 18M. **c)** Quantification of poroelastic properties measured by the nanorheometric test, including maximum phase angle (*δ*_m_), self-stiffening ratio (*E*_H_/*E*_L_), hydraulic permeability (*k*), and isotropic nonfibrillar matrix modulus (*E*_m_) derived from the fibril-reinforced poroelasticity model (*n* = 5). **d)** Maximum pore pressure calculated from the fibril-reinforced poroelastic finite element model simulation at the frequency corresponding to *δ*_m_. Panels a, c: mean ± 95% CI, *: *p* < 0.05; n.s.: not significant. Each data point represents the average value of > 10 locations measured from one animal. A complete list of quantitative and statistical outcomes is summarized in Supplementary Table S3.

Because cartilage function depends not only on elastic modulus but also its ability to dissipate energy through fluid flow-induced water-matrix interactions,^48^ we next assessed the dynamic poroelastic properties using our custom AFM-nanorheometric test. We quantified the frequency-dependent complex modulus, |*E*\*|, and phase angle, *δ*, a measure of relative energy dissipation, across four decades of frequencies (1-1,000 Hz) (Fig. 2b). From these spectra, we calculated the self-stiffening ratio, *E*_H_*/E*_L_, representing the capacity of cartilage to increase its dynamic modulus in response to high frequency loading activities,^49^ and extracted tissue hydraulic permeability, *k*, using the fibril-reinforced poroelastic finite element model.^50^

At 9 months, despite the absence of changes in *E*_ind_ and *E*_H_/*E*_L_, *Dcn^cKO^* cartilage exhibited a 1.9 ± 1.3-fold (mean ± 95% CI) increase in *k* and a corresponding reduction in fluid pressurization (Fig. 2c,d and Extended Data Fig. 4), indicating a loss of cartilage energy dissipative function. Notably, poroelastic parameters were more sensitive to matrix disruption than quasi-static modulus measurements, revealing functional compromise that was not captured by indentation modulus alone. Although increased fibril alignment may transiently preserve elastic resistance, the altered matrix exhibited reduced aggrecan-dependent fixed charge density and impaired fluid pressurization. By 18 months, *Dcn^cKO^* cartilage demonstrated higher dynamic moduli, consistent with the fibrotic matrix deposition. However, this apparent increase in modulus was accompanied by a 2.1 ± 0.9 fold reduction in self-stiffening ratio, a 6.5 ± 4.7-fold increase in *k*, and diminished pore pressure (Fig. 2c,d and Extended Data Fig. 4). These findings illustrate a substantial loss of poroelastic energy dissipative function in this remodeled, fibrous-like aged tissue. Thus, despite increased modulus, the tissue loses the fluid-supported load distribution and energy dissipative properties characteristic of healthy hyaline cartilage. Collectively, these findings demonstrate that decorin is required to preserve both elastic and poroelastic energy dissipative functions of articular cartilage during aging. Importantly, functional impairment of matrix fluid pressurization precedes overt structural erosion.

### Early Matrix Destabilization Precedes Aberrant Transcriptional Changes in Resident Chondrocytes

Given the multifaceted roles of decorin in both matrix assembly and cellular function,^51^ we next investigated whether decorin loss directly altered chondrocyte fate or transcriptional programs during aging. We performed single-cell RNA sequencing (scRNA-seq) of articular cartilage using the 10x Genomics platform following established procedures.^52^ After quality control filtering and cell type annotation (Extended Data Fig. 5), matrix-producing cells from 9-month-old cartilage were clustered into five transcriptionally distinct sub-populations (Fig. 3a,b). The relative abundance and distribution of these clusters were comparable between *Dcn^cKO^*and control cartilage (Fig. 3c), illustrating preserved cellular composition and heterogeneity despite pronounced surface matrix remodeling. Among these five clusters, Cluster 4 corresponded to articular chondrocytes, characterized by high expressions of canonical markers *Col2a1*, *Acan*, *Sox9*, *Col11a1*, and *Col11a2* (Fig. 3b). *Dcn* expression was selectively reduced in this cluster of *Dcn^cKO^* cartilage (Fig. 3d-f), reaffirming the chondrocyte-specific ablation in the *Dcn^f/f^/AcanCre^ER^*model. In contrast, other clusters did not exhibit appreciable changes in the expression of *Dcn* or other matrix genes, further validating target specificity. Strikingly, transcriptional alterations within articular chondrocytes were minimal at this stage. Only seven differentially expressed genes (DEGs) were identified in Cluster 4 (aside from *Dcn*), and none were directly associated with ECM turnover (Fig. 3e). Expression levels of major anabolic (*Col2a1*, *Acan*), catabolic (*Mmp3*, *Mmp13*, *Adamts5*), and transcription factor (*Sox9*) genes were unchanged (Fig. 3f). In support, RNAscope analysis corroborated comparable *Col2a1* and *Acan* transcript levels between genotypes (Fig. 3g). At the protein level, however, *Dcn^cKO^* cartilage displayed reduced aggrecan content despite similar *Acan* transcriptional activity, while it maintained similar collagen II levels compared to the control (Fig. 3h). This dissociation between transcription stability and matrix protein loss supports a primary role of decorin in extracellular proteostasis rather than direct regulation of chondrocyte biosynthesis. Importantly, no induction of collagen I was detected at either the transcriptional or protein levels (Fig. 3f-h), indicating that the early surface fibrillar remodeling (Fig. 1g-l) was not driven by chondrocyte dedifferentiation or fibrotic reprogramming, but through collagen II structural remodeling. Collectively, these data show that early structural matrix disruption occurs independently of major changes in chondrocyte identity or transcriptional state.

**Figure 3.**
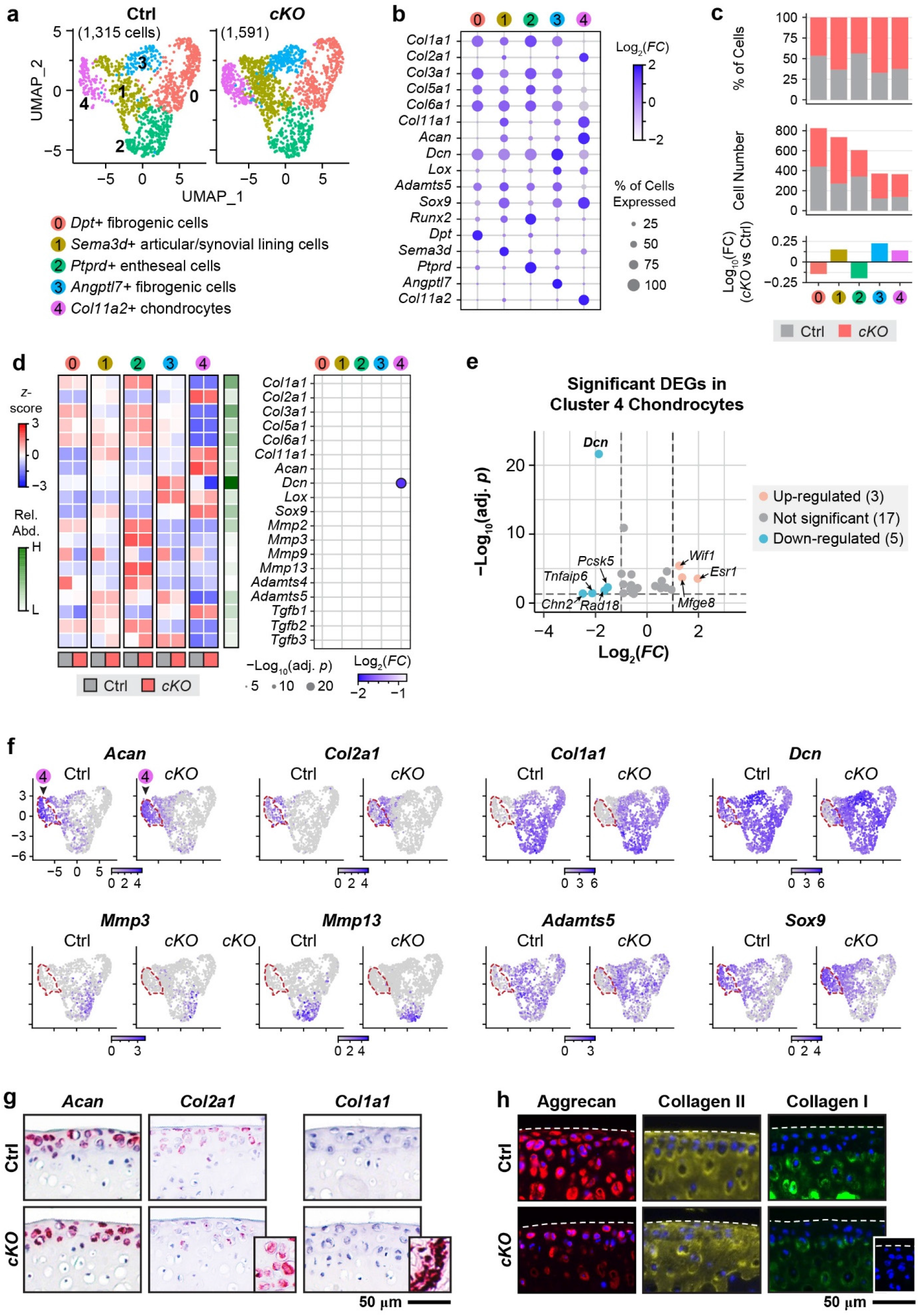
*Dcn^cKO^*cartilage exhibits limited alterations in cellular composition and transcriptional programming at 9 months of age. **a)** UMAP projections of control (Ctrl) and *cKO* cartilage cells, after excluding immune and hematopoietic cells. Clusters were annotated based on cluster-enriched marker genes and established lineage markers (see Extended Data Fig. 5). Cluster 4 corresponds to articular chondrocytes (*Acan*+, *Col2a1*+, *Col11a2*+, *Col1a1–*). **b)** Dot plot of representative ECM genes and top unique markers for each cluster. Dot size corresponds to the percentage of expressing cells; color denotes average log_2_ fold change (log_2_*FC*) relative to all other clusters. **c)** Relative proportion and absolute number of cells per cluster in Ctrl and *cKO* cartilage. Log_10_ fold change (Log_10_*FC*) of cell numbers represents differences in cluster abundance between genotypes. **d)** Left panel, heatmap of selected matrix anabolism, catabolism and TGF-β genes across clusters, row-scaled (*z*-score) to highlight relative expression differences between clusters and genotypes. The bar on the right shows relative expression abundance of each gene. Right panel, dot plot of significant differentially expressed genes (DEGs) of *cKO* versus Ctrl cells (|log_2_*FC*| > 1, adjusted *p* < 0.05). **e)** Volcano plot of significant DEGs in articular chondrocytes (Cluster 4). *Dcn* is significantly reduced in *cKO* chondrocytes. **f)** UMAP feature plots show the expression of representative cartilage ECM and catabolic genes in Ctrl and *cKO* cells. Cluster 4 cells are highlighted by pink dashed line, reaffirming the targeted reduction of *Dcn* in *cKO* articular chondrocytes (pink arrowhead). **g)** RNAscope *in situ* hybridization and **h)** immunofluorescence (IF) images show the spatial distributions of mRNA transcripts and proteins for *Acan*/aggrecan, *Col2a1*/collagen II and *Col1a1*/collagen I in femoral condyle cartilage (*n* = 4). Insets show the positive controls for RNAscope (*Col2a1*: meniscal horn, *Col1a1*: synovium) and internal negative control for IF.

By 18 months, in alignment with pronounced degenerative changes, scRNA-seq revealed broader cellular and transcriptional alterations in *Dcn^cKO^*cartilage. In addition to sharing clusters corresponding to chondrocytes, fibrogenic cells and synovial populations observed in controls, *Dcn^cKO^*cartilage contained a distinct additional population, Cluster 2, comprising approximately 20% of the matrix-producing cells (Fig. 4a-c and Extended Data Fig. 6). Within the articular chondrocyte cluster (Cluster 3), core matrix gene expression remained largely unchanged (Fig. 4d). However, the number of DEGs increased substantially compared to 9 months (Fig. 4e), with gene set enrichment analysis (GSEA) showing upregulated pathways related to TGF-β signaling, collagen fibril organization, and growth factor responses in *Dcn^cKO^* compared to control cartilage (Fig. 4f and Extended Data Fig. 7a). These data suggest perturbed signaling and cell-matrix interaction pathways in aged *Dcn^cKO^* cartilage, despite relative preservation of major anabolic gene expression. Meanwhile, other clusters also exhibited altered expression of collagens (Fig. 4d), consistent with active remodeling (Fig. 1). Despite this active remodeling, for articular chondrocytes at both ages, evaluating chondrocyte senescence burden using the *SenMayo* module score,^53^ we did not find significant differences in *SenMayo* scores or the expressions of canonical senescence markers *Cdkn2a* (p16^INK4a^), *Cdkn1a* (p21^CIP1^), and *Trp53* (p53)^8,9^ between genotypes (Extended Data Fig. 8), suggesting that the progressive matrix remodeling in *Dcn^cKO^* cartilage were not directly driven by chondrocyte senescence.

**Figure 4.**
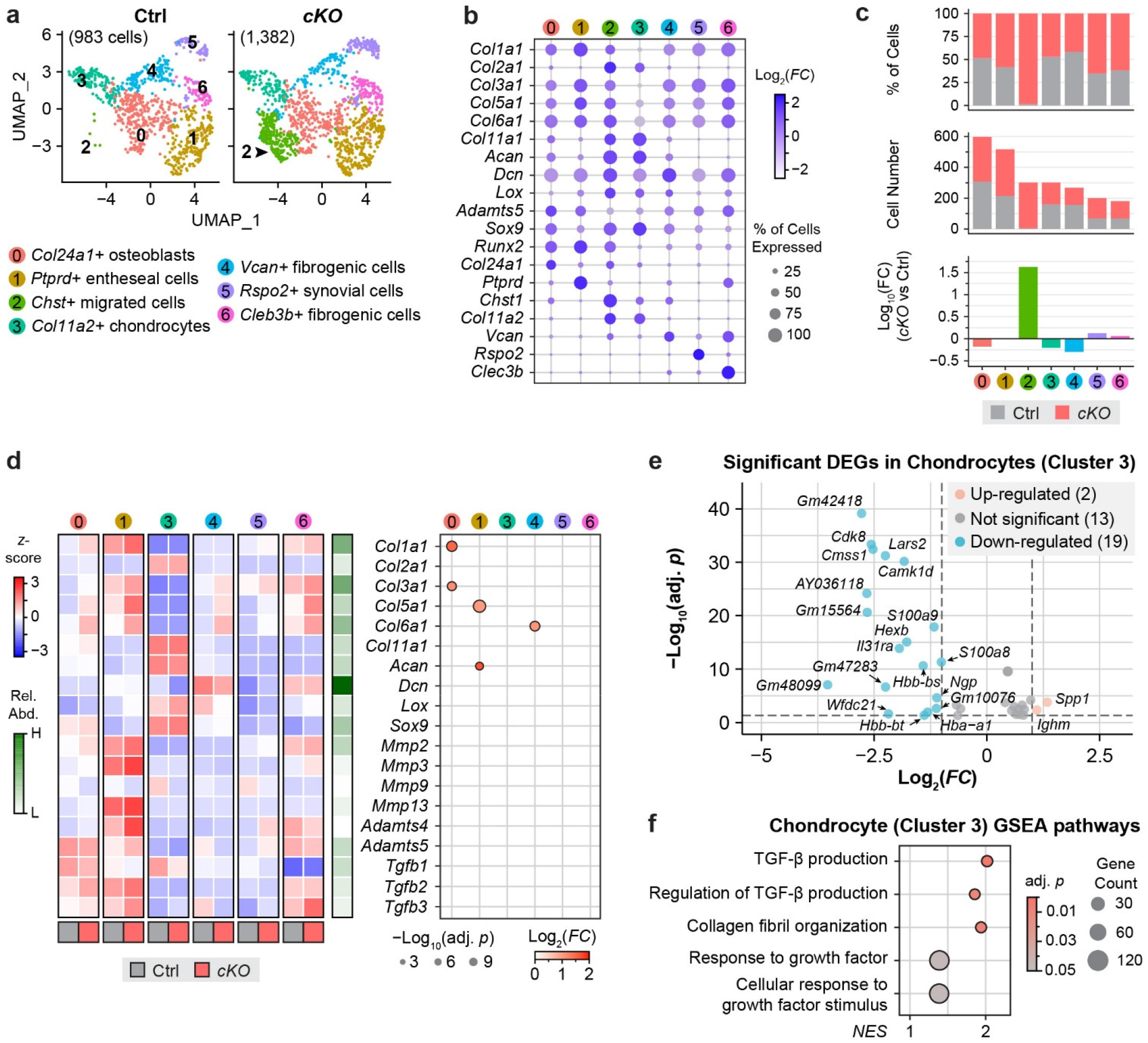
*Dcn^cKO^* cartilage develops altered cellular composition and transcriptional activities at 18 months of age. **a)** UMAP projections of control (Ctrl) and *cKO* cartilage cells, after excluding immune and hematopoietic cells. Clusters were annotated based on cluster-enriched marker genes and established lineage markers (see Extended Data Fig. 6). Cluster 3 corresponds to articular chondrocytes (*Acan*+, *Col2a1*+, *Col11a2*+, *Col1a1–*). Arrow highlights the emergence of Cluster 2 celles in *cKO* tissues. **b)** Dot plot of representative ECM genes and top unique markers for each cluster. Dot size corresponds to the percentage of expressing cells; color denotes average log_2_ fold change (log_2_*FC*) relative to all other clusters. **c)** Relative proportion and absolute number of cells per cluster in Ctrl and *cKO* joints. Log_10_ fold change (Log_10_*FC*) of cell numbers represents differences in cluster abundance between genotypes. **d)** Left panel, heatmap of selected matrix anabolism, catabolism and TGF-β genes across clusters, row-scaled (*z*-score) to highlight relative expression differences between clusters and genotypes. The bar on the right shows relative expression abundance of each gene. Right panel, dot plot of significant differentially expressed genes (DEGs) of *cKO* versus Ctrl cells (|log_2_*FC*| > 1, adjusted *p* < 0.05). **e)** Volcano plot of significant DEGs in articular chondrocytes (Cluster 3). **f)** Gene set enrichment analysis (GSEA) of Cluster 3 DEGs identifies pathways associated with TGF-β signaling, collagen fibril organization, and growth factor responses. A complete list of the top 20 positively and negatively enriched pathways (adjusted *p* < 0.05) is provided in Extended Data Fig. 7a.

The emergent Cluster 2 population in *Dcn^cKO^* cartilage displayed high metabolic and matrix-producing activities, marked by elevated expression of *Col1a1*, *Col2a1*, *Acan*, and *Dcn* (Fig. 5a). Its transcriptional profile was distinct from resident chondrocytes, as shown by the top marker genes (Fig. 5b and Extended Data Fig. 7b). CellChat cell-cell communication analysis revealed strong outgoing signals involving *Thbs*, *Mif*, *Fgf*, and *Jam* pathways (Extended Data Fig. 9), suggesting active matrix remodeling and immune-related interactions. Compared to articular chondrocytes, Cluster 2 cells exhibited enrichment of gene programs associated with ECM remodeling, cell adhesion, and positive regulation of skeletal development (Fig. 5c,d). Spatial mapping of the cluster marker *F5* confirmed localization of these cells within the “scar-like” fibrotic layer covering the eroded femoral surface (Fig. 5e). These cells expressed high levels of decorin (Fig. 5a,e,f), suggesting that they had not originated from *AcanCre^ER^*-targeted resident chondrocytes. Instead, their transcriptional profile and localization suggest infiltration from adjacent tissues. We found that in murine OA anterior cruciate ligament rupture (ACLR) OA model,^52^ the synovium contained an osteochondral progenitor population that shared similar top marker genes as these Cluster 2 cells (Extended Data Fig. 10), supporting that these cells were likely to be migrated from adjacent synovium tissues. Thus, early matrix destabilization precedes alterations in chondrocyte identity, whereas progressive architectural failure ultimately triggers aberrant signaling and recruitment of non-resident cell populations. Integrating findings across both timepoints supports a model in which matrix destabilization precedes and drives secondary cellular remodeling during age-associated cartilage degeneration.

**Figure 5.**
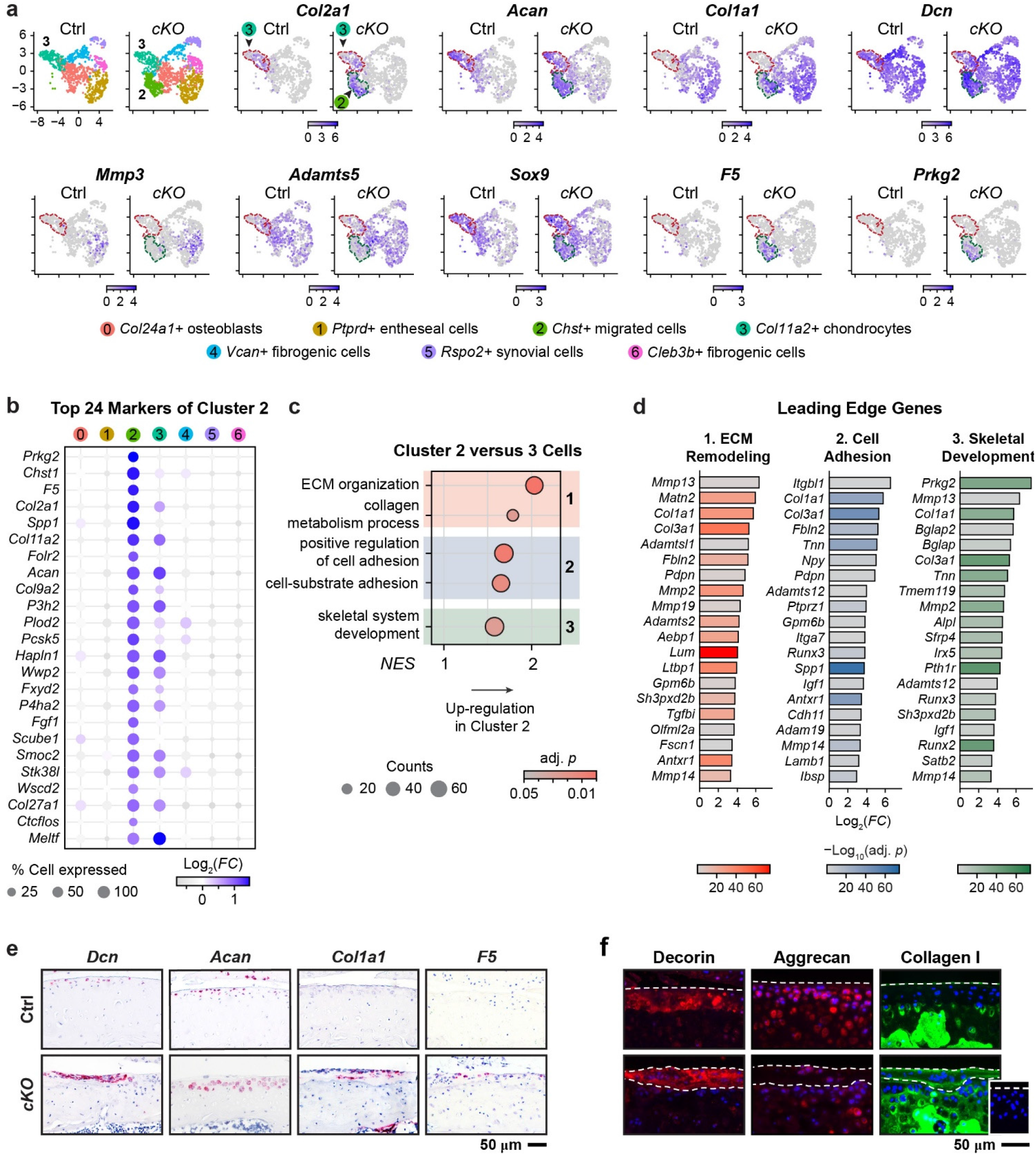
Transcriptional activities of the emergent cell population in *Dcn^cKO^* cartilage at 18 months. **a)** UMAP feature plots showing expression of representative ECM and remodeling genes in control (Ctrl) and *cKO* cartilage. Cluster 2 migrated cells and Cluster 3 articles chondrocytes are highlighted by dashed circles with colors corresponding to the original UMAPs. **b)** Dot plot of the top 24 Cluster 2 marker genes identified by *FindAllMarkers*. Dot size denotes the percentage of expressing cells; color corresponds to average log_2_ fold change (log_2_*FC*) for each gene within a given cluster relative to all other clusters. **c)** Gene set enrichment analysis (GSEA) comparing Cluster 2 versus 3 cells. Enriched pathways include ECM remodeling, collagen organization, cell adhesion, and skeletal system development. A complete list of positively and negatively enriched pathways (adjusted *p* < 0.05) is provided in Extended Data Fig. 7b. **d)** Top leading-edge genes contributing to enriched pathways associated with ECM remodeling, cell adhesion, and skeletal system development. **e)** RNAscope *in situ* hybridization images of *Dcn*, *Acan*, and *Col1a1*. Positive expression of *F5* is observed in cells within the scar-like fibrotic layer, consistent with Cluster 2 transcriptional signatures (*n* = 4). **f)** Immunofluorescence (IF) images of decorin, aggrecan, and collagen I. Insets show internal negative control (dashed white lines highlight cartilage surface and bottom edge of the fibrotic layer in *cKO* joints, *n* = 4).

### Decorin Directly Attenuates Collagen II Fibril Reorganization under Mechanical Load

Having established that early structural remodeling occurs independently of major transcriptional changes in resident chondrocytes, we then investigated whether decorin directly regulates collagen II fibril assembly and force-induced fibril organization. First, to assess the effects of decorin on individual collagen II fibrils, we induced fibrillogenesis of collagen II solution (0.3 mg/mL) within a PDMS microfluidic channel under controlled shear flow (19.2 µL/min) at 37°C (Fig. 6a).^54^ In the absence of decorin, self-assembled collagen II fibrils aligned robustly along the shear flow direction, as visualized by SEM. Addition of decorin at collagen II:decorin weight ratios of 25:1 and 15:1 significantly reduced fibril alignment (Fig. 6b,c), as quantified by the lower von Mises concentration, *κ* (Fig. 6d). Decorin also reduced fibril length and diameter (Fig. 6e), consistent with its role in modulating collagen fibrillogenesis. These findings demonstrate that decorin directly attenuates force-induced collagen II fibril alignment and growth.

**Figure 6.**
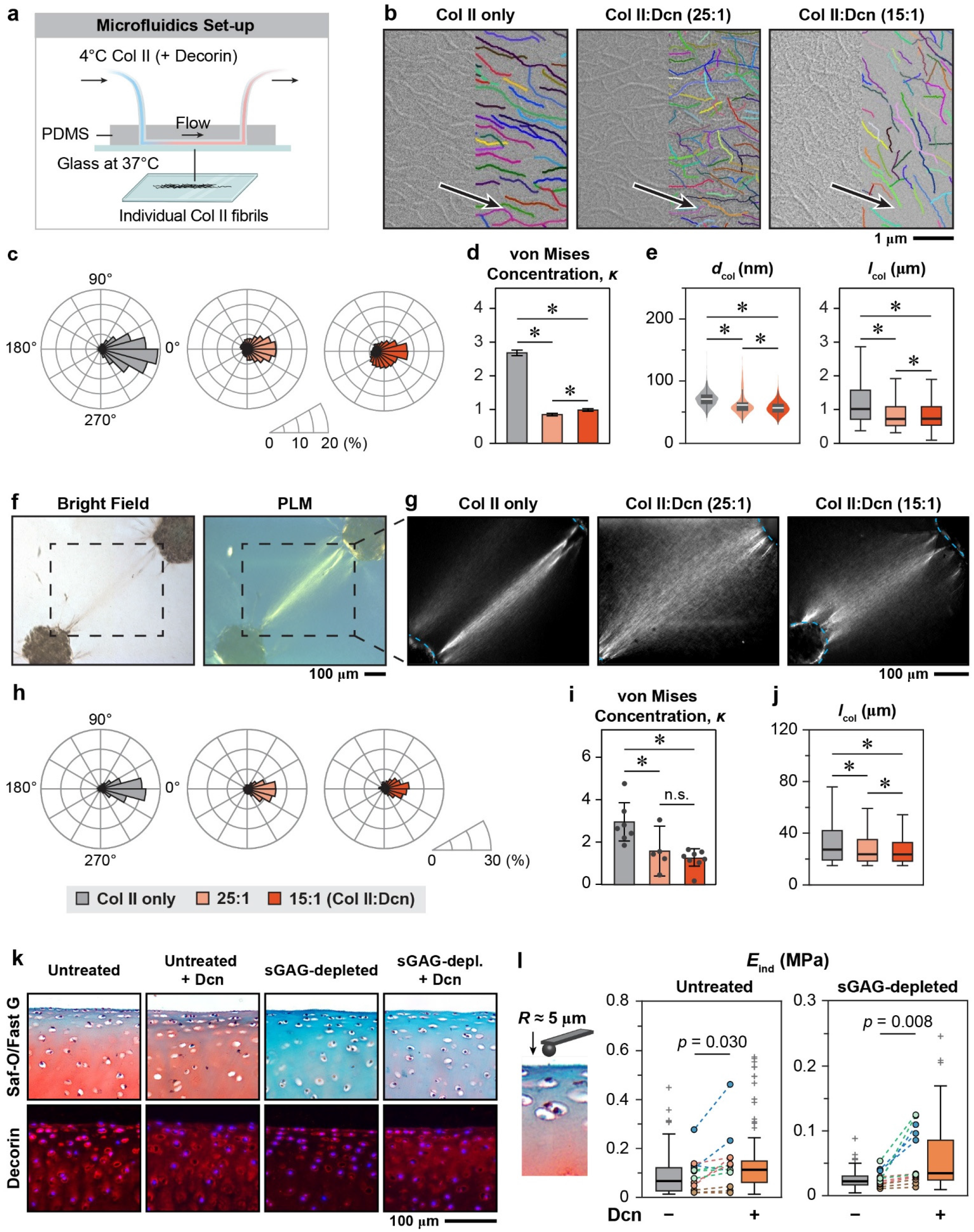
Decorin directly regulates collagen II fibril remodeling and strengthens cartilage surface collagen II fibrillar network. **a-e)** Decorin regulates the alignment of individual collagen II fibrils under fluid flow-induced shear stress. a) Schematic of the microfluidic setup for collagen II fibrillogenesis under shear flow. Cold bovine collagen II solution (with or without decorin) was polymerized within a PDMS channel at 37°C under a constant flow (19.2 μL/min) and deposited onto a glass substrate. b) Scanning electron microscopy (SEM) images of collagen II fibrils formed under shear flow in the absence or presence of decorin at collagen II:decorin ratios of 25:1 and 15:1 (w/w). Arrows denote flow direction. Overlaid lines represent tracking of individual fibrils by CT-FIRE. c) Rose plots of fibril orientation distribution, where 0° corresponds to the fluid flow direction. d) Fibril alignment quantified by the von Mises concentration parameter (κ, mean ± 95% CI). e) Violin and box-and-whisker plots of fibril diameter (*d*_col_) and length (*l*_col_) distributions measured from SEM images, respectively. Data in panels c-e represent > 4,500 fibrils pooled from three independent experiments for each condition (*: *p* < 0.05). **f-j)** Decorin mediates cellular tension-induced realignment of collagen II fibrils. f) Bright-field and polarized light microscopy (PLM) images of paired bovine chondrocyte spheroids seeded on collagen II hydrogels. g) PLM images showing collagen fiber compaction and alignment induced by spheroid contraction, which is reduced by the addition of decorin at 25:1 and 15:1 collagen II:decorin w/w ratios. Dashed lines denote spheroid boundaries. h) Rose plots of fiber orientation distribution from spheroid contraction assays, where 0° corresponds to the line of two spheroid centers. i) Fiber alignment quantified by κ (mean ± 95% CI). j) Fiber length (*l*_col_) derived from CT-FIRE analysis. Data in panels f and j represent ≥ 950 fibers pooled from ≥ 5 spheroid pairs across three independent experiments (*: *p* < 0.05; n.s., not significant). **k-l)** Decorin strengthens the collagen II fibrillar network on porcine cartilage surface. k) Safranin-O/Fast Green and decorin immunofluorescence (IF) images of untreated and sGAG-depleted porcine femoral condylar cartilage with or without decorin infiltration. l) Effective indentation modulus (*E*_ind_) on the cartilage surface with or without the infiltration of decorin measured by AFM-nanoindentation. Box-and-whisker plots show distributions of indentation values (10-15 locations per half-plug). Circles denote the mean value per half-plug. Dotted lines connect matched halves from the same plug. Different colors indicate individual animals (*n* = 4 animals, three plugs per animal).

To query this effect at the cellular scale, we next examined whether decorin regulates collagen II fibril organization under cell-generated mechanical forces. Using a modified spheroid contraction assay,^55^ two juvenile bovine chondrocyte spheroids were placed approximately 500 µm apart on 3 mg/mL collagen II hydrogels (Fig. 6f). Over 48 hours, cellular contraction generated long-range tensile forces that reorganized collagen II fibrils to a more aligned and compacted configuration between the spheroids, as illustrated by increased birefringence under polarized light microscopy (Fig. 6f, g). Incorporation of decorin at 25:1 and 15:1 collagen II:decorin ratios in the gel prior to spheroid seeding significantly reduced fibril alignment and compaction, reflected by decreased *κ* and reduced effective fibril length, *I*_col_, extracted from CTFire analysis (Fig. 6h-j). These data support that decorin incorporation into a type II collagen fibril network actively resists cellular force-mediated reorganization.

Finally, to determine whether decorin directly strengthens the native collagen II fibril network on cartilage surface, we performed AFM-nanoindentation on the surface of porcine femoral condylar cartilage, limiting the indentation-induced strain to the top ≈ 10 µm layer of the superficial zone. For intact cartilage, infiltration of decorin (20 µg/mL) significantly increased the surface indentation modulus by 53 ± 44% (mean ± 95% CI, Fig. 6k,l), illustrating reinforcement of the superficial matrix layer. To isolate effects on the fibrillar network, we enzymatically depleted sGAGs and non-fibrillar components prior to decorin treatment. In this collagen II fibril-dominated matrix, decorin supplementation significantly increased surface indentation modulus by 129 ± 89% (Fig. 6k,l), supporting a strengthening effect of decorin on the collagen architecture of cartilage surface. Collectively, these results demonstrate that decorin directly modulates collagen II remodeling and realignment under external forces, and reinforces the cartilage superficial layer fibrillar matrix. These mechanistic findings provide a biophysical basis for the progressive architectural destabilization observed in our aging *Dcn^cKO^*cartilage model.

## DISCUSSION

This study identifies that age-associated cartilage degeneration arises, at least in large part, from progressive failure of extracellular matrix stabilization. Using post-maturation deletion of decorin as a targeted perturbation, we show that disruption of extracellular structural proteostasis initiates surface fibrillar remodeling and poroelastic dysfunction, preceding substantial transcriptional reprogramming of resident chondrocytes and driving secondary cell-mediated remodeling (Fig. 7). These findings support a matrix-first model of cartilage aging and contrast with prevailing cell-centric paradigms of aging that emphasize senescence, mitochondrial dysfunction, or chronic inflammation as primary triggers of tissue decline.^56^ In matrix-dominant tissues such as cartilage, cellular turnover is limited and extracellular components must endure decades of mechanical stress (e.g., more than 100 million loading cycles for cartilage within a 60-year human lifespan^20^). Because disruption of this structural maintenance program is sufficient to initiate functional decline, our work establishes extracellular structural proteostasis as a distinct and essential dimension of aging biology that governs the long-term durability of load-bearing tissues. This framework extends the concept of proteostasis, traditionally defined at the intracellular level, to the extracellular environment.

**Figure 7.**
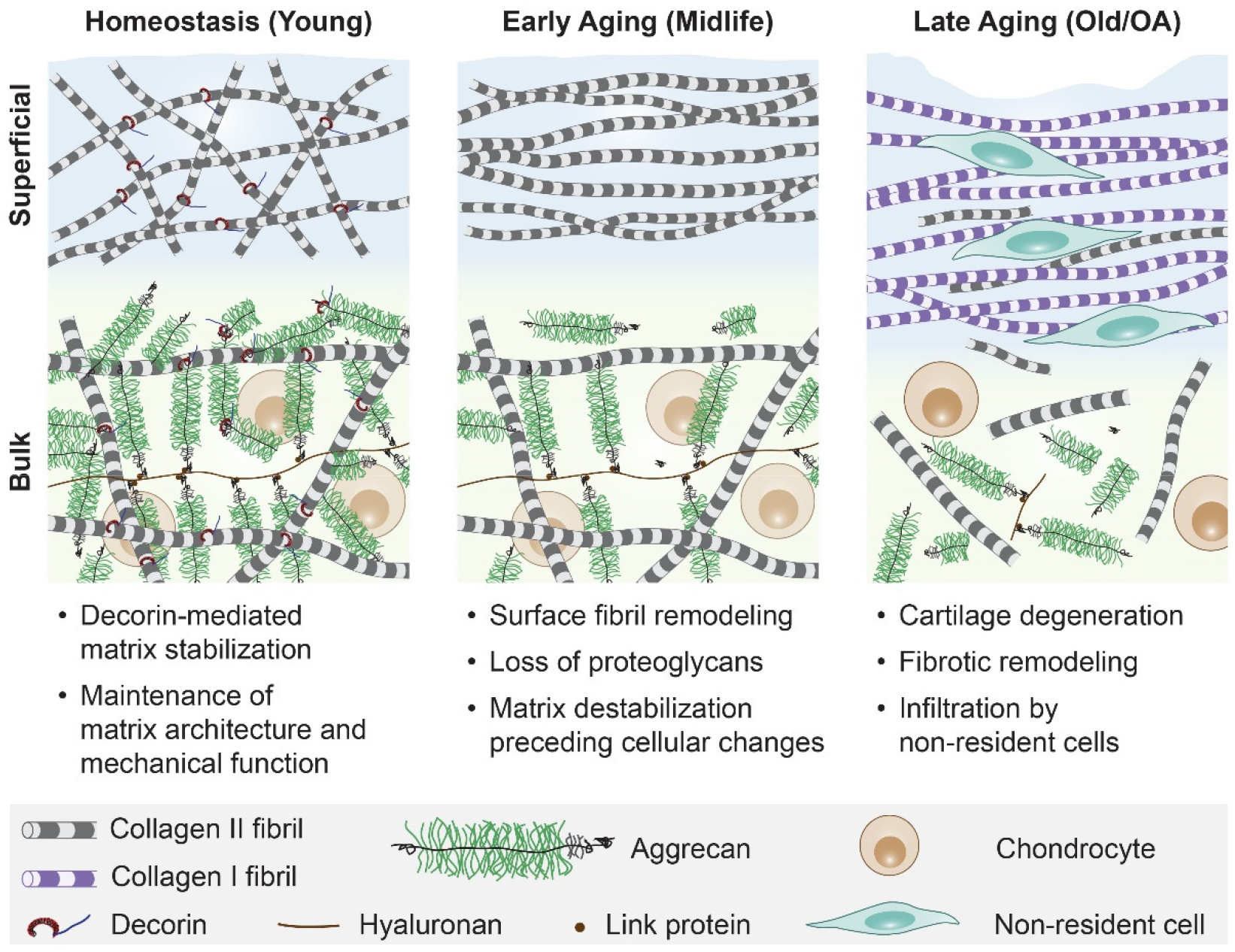
Matrix-first model of cartilage aging driven by failure of extracellular structural proteostasis. In young, normal cartilage, decorin stabilizes the superficial collagen II fibrillar network and promotes aggrecan retention within the matrix bulk, maintaining extracellular structural proteostasis and proper biomechanical functions. With the loss of decorin, during early aging, cartilage undergoes collagen fibril realignment and reduced aggrecan retention, resulting in accelerated surface fibril remodeling and impaired poroelastic mechanical properties, despite minimal transcriptional changes in resident chondrocytes. With advancing age, long-term matrix destabilization drives secondary cellular responses, including fibrotic remodeling, infiltration of non-resident cell populations, and cartilage erosion, culminating in osteoarthritis. These findings establish extracellular structural proteostasis as a primary determinant of cartilage integrity during aging.

A principal and previously unrecognized function revealed here is the role of decorin in maintaining the integrity of the superficial collagen II fibrillar architecture. The superficial zone of articular cartilage consists of a proteoglycan-poor, transversely isotropic meshwork of thin collagen II fibrils aligned tangentially to the surface.^47^ This specialized architecture resists shear forces,^57^ redistributes compressive stress to deeper layers,^58^ and supports interstitial fluid pressurization and lubrication.^59,60^ Disruption of this architecture is an early hallmark of osteoarthritic degeneration,^61,62^ leading to reduced tensile properties,^63–66^ increased friction,^67^ and elevated stresses within the cartilage bulk.^68^ We show that this fibrillar architecture depends on active stabilization by decorin during aging (Fig. 7). In *Dcn^cKO^* cartilage, aberrant fibril alignment and thickening emerge prior to overt cartilage erosion (Fig. 1i-l), supporting that decorin restrains force-induced remodeling of the superficial network under long-term joint loading. Importantly, this phenotype is not apparent in young adult *Dcn^−/−^* cartilage,^39^ suggesting that this architectural role of decorin becomes critical during prolonged mechanical use and aging rather than during development and maturation.

Mechanistically, stabilization of this fibrillar architecture is achieved through direct interactions between decorin and collagen II fibrils. Decorin binds to the fibril surface at multiple sites,^69^ and through its CS/DS-GAG chain, generates electrical double layer repulsion that limits lateral fusion and excess alignment of fibrils under shear and tensile stresses (Fig. 6). At the supramolecular level, decorin thereby prevents aggregation into aligned fibril bundles and preserves the transverse randomness characteristic of healthy cartilage surface (Fig. 1). Decorin dimerization^70^ may further contribute to interfibrillar linkage, reinforcing the network resistance to mechanically induced remodeling. Loss of decorin thus promotes fibril thickening and alignment along the split-line direction, where tensile stresses are greatest,^71^ consistent with our in vivo observations (Fig. 1) and in vitro biophysical assays (Fig. 6).

In addition to this architectural role, our findings reaffirm that decorin stabilizes aggrecan retention within the matrix bulk during aging. Consistent with our previous studies of postnatal maturation^39^ and post-traumatic degeneration,^41^ *Dcn^cKO^* cartilage exhibits reduced aggrecan content and impaired poroelastic fluid pressurization, despite the preserved *Acan* transcriptional activity of resident chondrocytes. These observations suggest that decorin-mediated aggrecan stabilization persists across the lifespan and remains essential for maintaining bulk matrix integrity. Thus, decorin exerts coordinated structural functions across all cartilage zones, safeguarding superficial fibril architecture while stabilizing collagen II-aggrecan integration within the matrix interior (Fig. 7). Together, these activities establish decorin as a central regulator of ECM proteostasis in aging cartilage.

The structural role of decorin in aging cartilage differs from its established biological activities in other contexts. Although decorin interacts with numerous growth factors and receptors,^51^ including TGF-β family members,^72^ the minimal early transcriptional changes observed following decorin ablation indicate that its predominant role in post-maturation cartilage maintenance is structural rather than signaling-mediated. Moreover, its regulation of collagen II fibril organization contrasts with its classical role in modulating lateral growth and tensile properties of collagen I matrices,^37^ highlighting the context-dependent nature of ECM regulation in vivo.

Collectively, these findings support a sequential model in which primary extracellular destabilization precedes and drives secondary cellular responses (Fig. 7). Early disruption of fibrillar organization and poroelastic function occurs in the absence of major transcriptional changes, supporting that initial degeneration is mechanically and structurally mediated. Our data show that decorin stabilizes the matrix by attenuating force-induced collagen II fibril realignment and preserving aggrecan-dependent fluid pressurization, thereby maintaining the mechanical integrity of cartilage under long-term loading. Loss of this stabilization leads to progressive architectural disorganization and impaired biomechanical functions. While chondrocytes initially tolerate moderate matrix perturbations, persistent destabilization ultimately disrupts mechanosensitive signaling, leading to aberrant pathway activation, recruitment of non-resident cell populations, and fibrotic remodeling. This temporal and mechanistic hierarchy positions ECM failure upstream of canonical cellular hallmarks of aging in this tissue context.

The matrix-first framework established here may extend broadly to other long-lived, mechanically loaded tissues with limited regenerative capacity, including intervertebral disc, tendon, and sclera. In these systems, extracellular components must withstand cumulative mechanical stress over decades, suggesting that progressive failure of matrix stabilization may represent a common initiating mechanism of age-associated degeneration. This perspective shifts the focus from cellular decline alone^56^ to the durability of extracellular architecture as a central determinant of tissue aging. Therefore, therapeutic strategies aimed at reinforcing matrix organization and stability may complement existing approaches targeting cellular rejuvenation, particularly in matrix-dominated tissues where regenerative capacity is limited.

This study demonstrates that age-associated cartilage degeneration arises, at least in large part, from progressive failure of extracellular matrix stabilization. By preserving the superficial collagen II fibrillar architecture and aggrecan-dependent poroelastic function, decorin acts as a structural safeguard under lifelong mechanical loading. Loss of this stabilization disrupts matrix integrity prior to major transcriptional reprogramming of resident chondrocytes, ultimately leading to tissue degeneration. These findings establish extracellular structural proteostasis as a primary determinant of tissue longevity and support a matrix-first framework for aging in load-bearing connective tissues. Targeting mechanisms that reinforce matrix stabilization may therefore provide a complementary therapeutic axis to mitigate age-associated degeneration in cartilage and other mechanically loaded tissues.

## METHODS

### Animal model

Cartilage-specific inducible decorin knockout mice (*Dcn^flox/flox^/AcanCre^ER^*, or *Dcn^cKO^*) in the C57BL/6 background were generated by breeding decorin-floxed (*Dcn^flox/flox^* or *Dcn^f/f^*) mice^73^ with mice containing a knock-in of the tamoxifen (TM) inducible Cre in the aggrecan gene (*AcanCre^ER^*,^74^ Jackson Laboratory). Tamoxifen was injected intraperitoneally to 3-month-old mice for 3 consecutive days at 4.5 mg/40 g body weight in the form of 20 mg/mL suspension in sesame oil (S3547, Sigma) with 1% volume/volume benzyl alcohol (305197, Sigma). Three age-matched control groups with normal expression of decorin were used, including *Dcn^flox/flox^/AcanCre^ER^*mice injected with vehicle (sesame oil and benzyl alcohol without tamoxifen), as well as *Dcn^flox/flox^* and wild-type (WT) mice injected with tamoxifen at the same dose and frequency. We found no differences amongst these three controls. Mice were euthanized at 4, 9, and 18 months of age to evaluate phenotypic changes. All the animals used were genotyped for *Dcn* and *AcanCre^ER^* by PCR of DNA extracted from tail biopsy specimens following standard procedures.^73^ Both male and female mice were included, as we did not observe sex-associated variations in the phenotype (Extended Data Fig. 1e-f).^39^ All animal work was approved by the Institutional Animal Care and Use Committee (IACUC) at Drexel University.

### Histology and immunofluorescence (IF) imaging

Whole knee joints were fixed in 4% paraformaldehyde (PFA) for 24 hours, decalcified in 10% EDTA for 21 days, and embedded in paraffin. Serial six μm-thick sagittal sections were prepared and stained with Safranin-O/Fast Green to evaluate gross-level joint morphology, sGAG staining, and cell density (*n* = 6). For each joint, approximately 15 sections were obtained and scored by two blinded observers (BK and YL) using the modified Mankin method.^75^ Also, synovial histopathology scores were assessed by two blinded observers (TL and SYK) to assess changes including lining hyperplasia, fibrosis, and sublining cellularity following standard procedures.^76^ Thicknesses of uncalcified and total cartilage were quantified by averaging six values evenly distributed across the entire cartilage in the load bearing medial region using ImageJ. Picrosirius Red staining (ab150681, Abcam) was applied to evaluate collagen fiber organization, with imaging performed on a Leica DMLP polarizing light microscope (PLM), following standard procedures.^77^

For IF imaging, after deparaffinization, sections were treated with 0.1% pepsin (P7000, Sigma) or 500 U/mL hyaluronidase (H3506, Sigma) for antigen retrieval for collagen I, collagen II and decorin. For aggrecan, antigen retrieval was done by first treating the sections with 10 μg/mL of proteinase K (P6556, Sigma) for 20 minutes at 37°C, following with 1.5 mg/mL hyaluronidase (H3506, Sigma) for 30 min at 37°C, then 14 hours incubation at 1× citrate buffer (HK0809K, Biogenex) at 65°C. Sections were then incubated with primary antibodies in 1% BSA and 5% goat serum (50062Z, Invitrogen) for collagen I (1310-01, 1:200 dilution, SouthernBiotech), collagen II (MAB8887, 1:200, Sigma), decorin (ENH019-FP, 1:100, Kerafast), and aggrecan (AB1031, Sigma) overnight at 4°C. Next, sections were incubated with corresponding secondary antibodies for collagen I (A11055, 1:500, Invitrogen), collagen II (A11001, 1:200, Invitrogen), decorin (A11008, 1:500, Invitrogen) and aggrecan (A11008, 1:200, Invitrogen), for 1 hour at room temperature, followed by DAPI Fluoromount-G (no. 0100-20, SouthernBiotech) mounting, and imaged with a Leica DMI-6000B microscope. Internal negative controls were prepared and imaged following the same procedure, except without primary antibody incubation.

### RNAscope in situ hybridization

RNAscope in situ hybridization was performed using the RNAscope2.5 HD Detection reagent-RED kit (322350, ACDbio) to assess the spatiotemporal expression of specific murine genes including *Col1a1* (319371, ACDbio), *Col2a1* (407221, ACDbio), *Acan* (439101, ACDbio), *Dcn* (413281, ACDbio), and *F5* (502411, ACDbio). Paraffin-embedded sections were deparaffinized through a graded xylene and ethanol series, followed by pretreatment with a custom reagent (300040, ACDbio) for 30 minutes at 40°C. Probe hybridization was conducted at 40°C for 2 hours in a temperature-controlled hybridization oven, followed by signal amplification and detection per the manufacturer’s protocol.^78^ For visualization, sections were counterstained with Hematoxylin (51257-100ml, Sigma), dried and sealed with Permount mounting medium (SP15-100, Fisher Scientific), and then, imaged using a Leica brightfield microscope.

### Scanning electron microscopy (SEM)

Freshly dissected femoral condyles were treated with 0.1% trypsin (T7409, Sigma) and 500 U/ml hyaluronidase (H3506, Sigma) at 37°C each for 24 hours to remove proteoglycans. Samples were then fixed with 4% PFA, dehydrated through graded water-ethanol and ethanol-hexamethyldisilazane (HMDS) mixtures, and air-dried overnight. SEM images were acquired using a Zeiss Supra 50VP microscope on samples coated with a 6-nm-thick platinum layer, following established procedures.^79^ Collagen fibril thickness, *d*_col_, and alignment angle, *θ*, were quantified using ImageJ (≥ 100 fibrils per animal, *n* = 4). Values of *θ* were multiplied by two to account for the absence of directionality^80^ and offset to the medial-lateral joint orientation as 0°. Pooled *θ* values were then fitted to the von Mises probability density function to derive the von Mises concentration, *κ*.^81^

### AFM-based nanomechanical tests

Classical AFM-based nanoindentation was applied to the medial side of freshly dissected femoral condyle cartilage (*n* ≥ 5 animals) using a polystyrene microspherical tip (*R* ≈ 5 μm, nominal *k* ≈ 8.9 N/m, HQ:NSC35/tipless/Cr-Au, NanoAndMore) and a Dimension Icon AFM (Bruker Nano) in 1× PBS containing protease inhibitors (Pierce 88266, Fisher Scientific).^39^ For each sample, at least 10-15 different locations were tested on the load-bearing region at a 10 μm/s *z*-piezo displacement rate. To ensure that cartilage deformation was restricted to the linear, small-deformation regime, the maximum force was limited to ≈ 2 µN, which corresponded to ∼ 1 µm maximum depth. The effective indentation modulus, *E*_ind_, was calculated by fitting the entire portion of *F-D* loading curve to the Hertz model, assuming a Poisson’s ratio *ν* ≈ 0.1.^39^

A custom-built AFM-nanorheometric test was applied to quantify the time-dependent poroelastic properties of femoral condyle cartilage (*n* ≥ 5), following established procedures.^39^ In brief, a microspherical tip (*R* ≈ 12.5 μm, nominal *k* ≈ 16 N/m, HQ:NSC35/tipless/Cr-Au, NanoAndMore) was programmed to indent into the sample up to a static depth of *D*_0_ ∼ 1 μm, resulting in an effective contact radius ≈ 5 μm. The tip was held at the constant position for 90 seconds to allow force relaxation, during which a random binary sequence (RBS) displacement with an amplitude of ≈2–3 nm was superimposed on the static depth, enabling the measurement of dynamic mechanical behaviors corresponding to 1−1000 Hz frequencies. The output force-displacement data were processed with discrete Fourier transform to extract the frequency components of the dynamic force, *F*\* and displacement, *D*\*. The magnitude of the dynamic complex indentation modulus, |*E*\*|, was calculated by the Taylor series expansion of the Hertz model, *F*^∗^ ≈ 2|*E*^∗^|*R*^1/2^*D*^1/2^*D*^∗^/(1 − *v*^2^).^82^ The phase angle, *δ*, was calculated as the phase lag of *D*\* relative to *F*\*. For each test, the low frequency modulus, *E*_L_, and high-frequency modulus, *E*_H_, were calculated as the average magnitude of the dynamic modulus, |*E*\*|, at ≤ 5 Hz and at 800−1200 Hz, respectively.

A fibril-reinforced poroelastic finite element model was applied to the spectra of |*E*\*| and *δ*, to calculate the hydraulic permeability, *k*.^50^ Cartilage was modeled as a composite of an isotropic nonfibrillar matrix, a fibril network and a fluid phase. The properties of the nonfibrillar matrix were described by the elastic modulus, *E*_m_, hydraulic permeability, *k*, and Poisson’s ratio *ν*. The fibril network was modeled with a tensile-only Young’s modulus, *E*_f_. The deformation and pore pressure fields caused by static (*D*_0_ ≈ 1 µm) and dynamic compression were confined to the top ≈ 10 µm range.

### Microcomputed tomography (μCT)

Micro-computed tomography (μCT) was applied to fixed knee joints using a MicroCT 35 (Scanco Medical). Scans were conducted at a 6 μm isotropic voxel size, and images were smoothed with a Gaussian filter (Sigma = 1.2, support = 2.0). Regions of interest (ROIs) for the subchondral bone plate (SBP) and subchondral trabecular bone (STB) were defined at a threshold of 30% of the maximum image gray scale. The SBP of the tibial plateau’s medial side central loading region was contoured to calculate SBP thickness (SBP.Th). For STB, the load-bearing ROI on the medial side was analyzed to quantify structural parameters, including bone volume fraction (BV/TV), trabecular number (Tb.N), thickness (Tb.Th), and separation (Tb.Sp). These measurements were performed using Scanco’s 3D standard microstructural analysis software, following established procedures.^42^

### Single-cell RNA sequencing

Single-cell RNA sequencing (scRNA-seq) was performed on femoral condyle and tibial plateau cartilage isolated from 9M and 18M mice (four joints pooled from one male and one female mice for each group, Table S7). Immediately after euthanasia, cartilage tissues were carefully peeled off with a blade, with excess bone, ligaments, and fat tissues removed. Harvested cartilage was washed twice in 1× PBS to remove tissue debris and blood cells. Cells were isolated using enzymatic digestion with pronase (53702-250KU, Millipore) and collagenase II (17101-015, Gibco) supplemented with DNase (DN25-100MG, Sigma) for 1 hour, following an established procedure.^83^ After digestion, a red blood cell removal kit (130-094-183, Miltenyi Biotec) and a dead cell removal magnet kit (130-090-101, Miltenyi Biotec) were applied on ice. The cell suspension was strained twice to ensure a high quality single-cell suspension. Single-cell suspensions with > 90% viability were submitted to the Center for Applied Genomics at the Children’s Hospitald of Philadelphia, and processed via the 10x Genomics pipeline (Chromium Next GEM Single Cell 3’ Kit v3.1) for barcoding and library preparation. Raw data (FASTQ files) and filtered barcodes, features, and matrix files were deposited in GEO under the file names listed in Table S7.

Seurat (R, v4.1.0) was employed for quality control and data analysis, following established procedures.^52^ Quality control, normalization, dimensionality reduction, and unsupervised clustering were performed as described in detail the Supplementary Methods. In brief, low-quality cells (including multiplex, debris, low viability cells) were excluded based on *nFeature*, *nCount*, and mitochondrial gene content thresholds. Cell cycle and mitochondrial gene expression variation were regressed out prior to scaling. Principal component analysis (PCA) followed by Uniform Manifold Approximation and Projection (UMAP) was used for dimensionality reduction, and distinct cell clusters were identified using *FindNeighbors()* and *FindClusters()*. Cluster marker genes were identified using *FindAllMarkers()*, retaining genes with positive differential expression and high cluster specificity. Differentially expressed genes (DEGs) between control and *cKO* conditions were identified using *FindMarkers()*. Gene Set Enrichment Analysis (GSEA) was performed using the *fgsea* R package on genes pre-ranked by log2 fold-change (Log_2_*FC*), and significantly enriched pathways (adjusted *p*-value < 0.05) were visualized as dot plots in *ggplot2*. Intercellular communication was inferred using *CellChat*, with communication probabilities computed for each ligand-receptor interaction and dominant incoming and outgoing signaling patterns identified via non-negative matrix factorization. Relationships between cell populations, communication patterns, and signaling pathways were visualized using alluvial plots. For the analysis of chondrocyte senescence, module scores for the SenMayo gene signature were computed using *AddModuleScore()* and visualized with *Featureplot*. The SenMayo mouse gene set comprises 118 senescence and SASP-associated genes;^53^ 108 and 109 genes were detected at 9M and 18M respectively (coverage: 91.5% and 92.4%). Additionally, normalized expressions of canonical senescence effectors *Cdkn2a*, *Cdkn1a*, and *Trp53* were extracted from the data, as well as the cell percentage expressing them. Comparisons were performed using the Mann-Whitney U test for mean expression and Fisher’s exact test for proportions of cell expressing the markers. For the analysis of 18-month-old groups, to assess the transcriptional identity of Cluster 2 migrated cells in the context of known synovial cell populations, the top 24 Cluster 2 marker genes were projected as a gene module score onto a published single-cell atlas of synovial fibroblasts comprising seven subpopulations,^52^ and expression of the leading marker *F5* was independently visualized across the same atlas.

### Collagen II fibrillogenesis under shear flow via a microfluidic system

A custom microfluidic system was applied to assess the impact of decorin on shear flow-induced collagen II fibril alignment. In brief, a polydimethylsiloxane (PDMS) cast mold was sealed to an oxidized glass coverslip, which were put on a hotplate and infused with a cold collagen II solution containing decorin at different ratios. Formation, alignment and deposition of collagen II fibrils on the glass substrate were induced by controlled fluid flow at 37°C. After infusion, the PDMS stamp was peeled off to enable SEM imaging of the assembled fibrils.^54^

### PDMS device casting

The glass coverslip substrate was cleaned in an ultrasonic bath with DI water to remove dust and then oxidized by boiling for 20 minutes in a mixture of water, 27% ammonium hydroxide, and 30% hydrogen peroxide at a 5:1:1 volume ratio at 70°C for immediate use. The slides were then rinsed in DI water and dried in air. PDMS devices were fabricated by pouring a degassed, well-mixed Sylgard 184 Silicone Elastomer (NC9285739; Fisher Scientific) with a curing agent at a 10:1 mass ratio over a plasma-coated silicon master fabricated by GeSiM Bioinstruments and Microfluidics.^84^ The mixture was allowed to cure for 2 hours in a desiccator at 60°C. The resulting PDMS mold contained microchannels measuring 8 mm in length, 500 µm in width, and 50 µm in height. The PDMS and coverslip were sealed together using an additional integrated channel and a vacuum pump. A syringe pump was connected through polyether-ether-ketone (PEEK) tubing to the channel plate (GeSiM Bioinstruments and Microfluidics) to enable the control of collagen solution flow rate in the microchannels (Fig. 6a).^54^

### Collagen solution and flow conditions

Bovine collagen II (C1188, Sigma) was dissolved in 10 mM acetic acid at 0.3 mg/mL, with osmolarity and pH adjusted to physiological conditions using 10× PBS, DI water, and NaOH. Bovine decorin (D8428, Sigma) was added to the collagen solution at concentrations of 0.012 mg/mL (25:1 collagen II:decorin w/w ratio) and 0.02 mg/mL (15:1 w/w ratio). To prevent premature fibrillogenesis, all components were mixed and kept in an ice bath before use. The syringe pump and connecting tubes were placed in a cold Styrofoam box at 4°C. Meanwhile, the PDMS device with the sealed coverslip was placed on a hotplate and heated to 37°C. The flow system was degassed and rinsed with 4 mL of cold DI water before introducing collagen solution. Next, the cold collagen II solution was drawn into chilled syringes and immediately pumped from the cold reservoir into the heated PDMS channels. The initial flow rate was set at 480 μL/min for approximately 50 μL of volume and stopped once the collagen solution reached the channel. The flow rate was then adjusted to 19.2 μL/min to allow the perfusion of collagen II + decorin solution through the microfluidic channels for 40 minutes. After perfusion, the channel was rinsed with DI water at 19.2 μL/min for 30 minutes to remove unattached collagen fibrils. Finally, the device was disassembled by peeling the PDMS stamps from the collagen fibril-coated glass coverslip. The coverslip was then dried on a hotplate at 37°C, coated with a 6-nm-thick platinum layer, and imaged using a Zeiss Supra 50VP SEM.

SEM images were analyzed with the CT-FIRE (Curvelet Transform-Fiber Extraction) program.^85^ Collagen fibril alignment (*θ*_col_) and thickness (*d*_col_) were quantified using MATLAB, analyzing 4500 fibrils pooled from three independent experiments per condition. Values of *θ*_col_ were multiplied by two to account for the lack of directionality^80^ and offset to the flow direction as 0°. The pooled *θ* data were fitted to the von Mises probability density function to calculate the concentration parameter, *κ*.^81^

### Collagen gel contraction assay

The collagen gel contraction assay was applied to assess the effects of decorin on chondrocyte-mediated collagen II fibril reorganization, following the procedure originally established for collagen I gels.^55^ Collagen II gels with different decorin concentrations were prepared in a 12-mm culture dish with glass bottom for chondrocyte spheroid seeding and culture. Bovine collagen II (C1188, Sigma) was dissolved in 10 mM acetic acid at 2.5 mg/mL with osmolarity and pH adjusted to physiological conditions using 10× PBS, DI water, and NaOH. Bovine decorin (D8428, Sigma) was added to the collagen solution at final concentrations of 0.1 mg/mL (25:1 collagen:decorin w/w ratio) and 0.17 mg/mL (15:1 w/w ratio). The collagen-decorin mixture was pipetted onto the culture dish and incubated at 37°C for 48 hours to allow gelation.

For chondrocyte isolation, Articular cartilage disks (diameter ≈ 5 mm, thickness ≈ 2 mm) were harvested from femoral condyles of ≈ 6-12-month-old calves (Green Village), digested with 125 U/mL collagenase I (17018029, Thermo Fisher) for 16 hours, and then, passed through a 70-μm strainer to remove undigested fragments. The filtered chondrocytes were used to form spheroids via the hanging droplet method.^86^ In brief, cells were suspended in chondrogenic DMEM at 25,000 cells/mL, and 20 μL droplets were placed on the underside of a petridish lid, with 10 mL 1× PBS added to the dish to maintain humidity. After 5 days of culture, spheroids were collected and carefully placed in pairs at ≈ 500 μm apart on the surface of collagen gels and cultured for an additional 48 hours. The gels were then fixed in PFA for 10 minutes and stained with Picrosirius Red (ab150681, Abcam) for 1 hour. Images were captured using Leica DMLP PLM and analyzed with CT-FIRE.^85^ Collagen fibril alignment (*θ*_col_) and length (*l*_col_) between spheroids were quantified using MATLAB. At least 1,000 fibrils from a minimum of five spheroid pairs across three independent experiments were analyzed per condition. Values of *θ*_col_ were multiply by two, offset to the line of two spheroid centers as 0°, and pooled *θ* data were fitted to the von Mises probability density function to calculate *κ*.^81^

### Porcine cartilage explant treatment and surface modulus quantification

Fresh articular cartilage disks containing intact surfaces (diameter ≈ 5 mm, thickness ≈ 2 mm) were harvested form ≈ 2-3-year old porcine knee joints (Green Village) using a biopsy punch. For GAG depletion, samples were first incubated with 0.5 U/mL chondroitinase ABC (MilliporeSigma, C3667) for 24 hours, followed by 500 U/mL hyaluronidase (MP Biomedial, 37326-33-3) in 1× PBS for an additional 24 hours. Untreated disks were maintained in 1× PBS with protease inhibitors for 48 hours. For both untreated and GAG-depleted disks, samples were bisected to generate matched halves. One half was incubated with bovine decorin (20 µg/mL in 1× PBS, D8428, MilliporeSigma) for 24 hours at 37°C, and the other half was incubated in 1× PBS. Classical AFM-nanoindentation was applied to the surfaces of both halved explants with a microspherical tip (*R* ≈ 5 μm, nominal *k* ≈ 5.4 N/m, HQ:NSC35/tipless/Cr-Au, NanoAndMore) and a Dimension Icon AFM (Bruker Nano) to quantify the indentation modulus, *E*_ind_. For each condition, experiments were repeated for 12 explants from four animals, and at least 10-15 locations were tested on each sample to account for spatial heterogeneity. By limiting the maximum indentation depth to be ∼ 1 µm, we ensured that the values of *E*_ind_ reflected the properties of top ∼ 10 µm superficial layer of porcine cartilage. In addition, Safranin-O/Fast Green and decorin IF imaging were applied to validate the sGAG depletion and decorin infiltration (*n* = 4 animals), respectively, as describe in previous sections.

### Statistical Analysis

Statistical tests were selected based on data characteristics, including sample size, distribution normality, continuity, and heteroscedasticity. For cartilage thickness, cell density, and modified Mankin scores, the non-parametric Mann-Whitney U test was performed to test the significance between genotypes. For murine cartilage biomechanical properties, *p*-values were calculated using a two-sample Welch’s *t*-test between genotypes. In addition, the effects of decorin on cartilage surface modulus were assessed by non-parametric Wilcoxon signed-rank test between paired explants. For nanostructural data, the fiber length (*l*_col_) and fibril diameter (*d*_col_) were compared between groups or genotypes using either unpaired two-sample *t*-test or one-way ANOVA followed by Tukey’s post hoc test. For von Mises concentration, *κ*, was compared between genotypes or groups via the Mardia-Jupp test,^87^ followed by Holm-Bonferroni correction for multiple comparisons. For all the tests, the significance level was set at *α* = 0.05. A complete list of all quantitative and statistical outcomes is summarized in Tables S1–S6.

## ACKNOWLEDGEMENTS

This work was financially supported by the National Institutes of Health (NIH) grants R01AR074490 (to LH) and P30AR096619 to the Penn Center for Musculoskeletal Disorders. Additional support was provided by the Department of Veterans Affairs grant IK6 RX003416 (to RLM) and the National Science Foundation (NSF) grant CMMI-1548571 to the Center for Engineering Mechanobiology.

## Data Availability Statement

The data that support the findings of this study are available from the corresponding author upon reasonable request.

## EXTENDED DATA

**Extended Data Figure 1.**
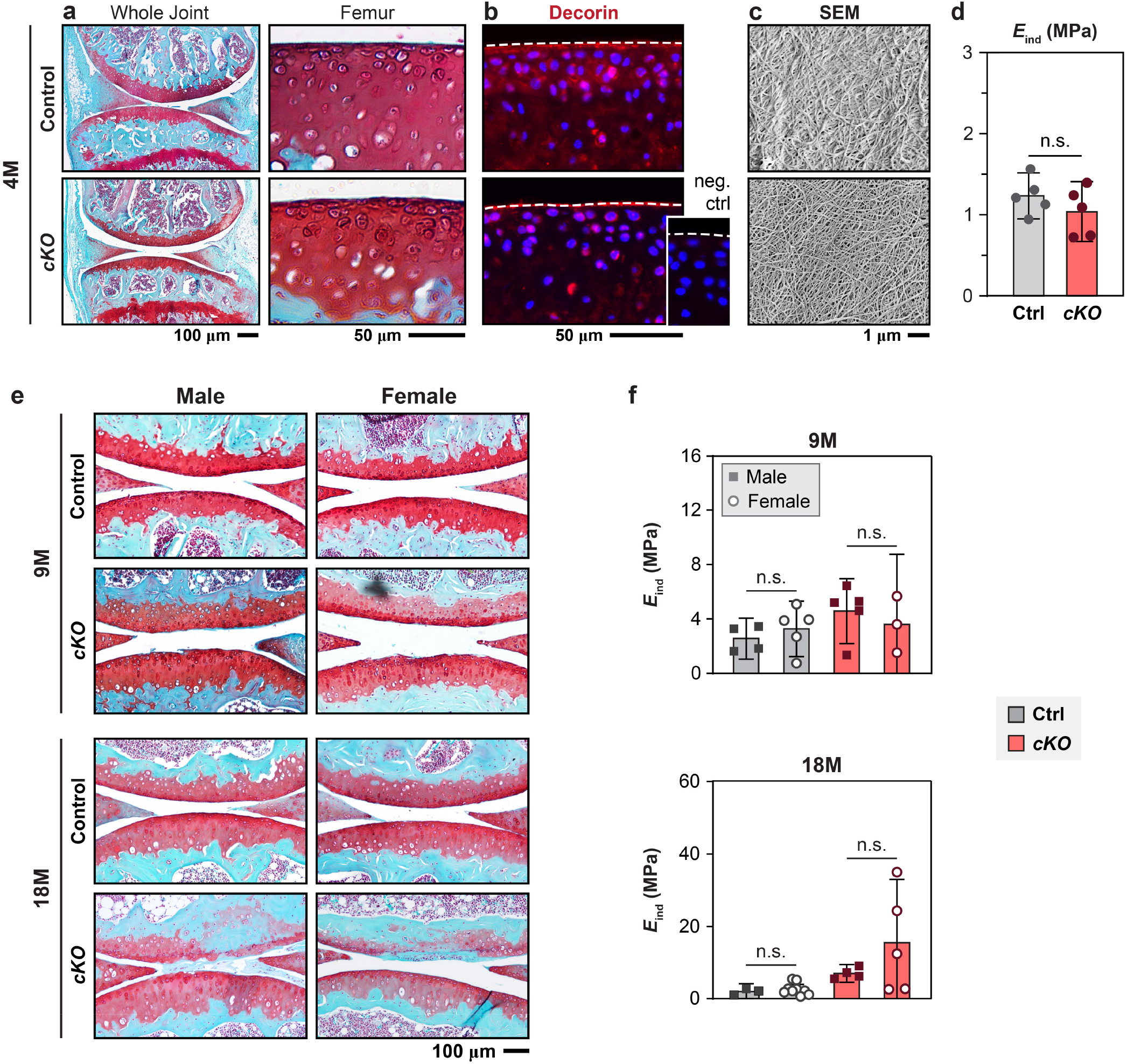
Validation of the absence of short-term or sex-dependent effects of decorin loss on cartilage integrity. **a-d)** Phenotypic changes of *Dcn^cKO^*cartilage at 4 months of age (4M). Representative a) Safranin-O/Fast Green histological staining, b) decorin immunofluorescence (IF), and c) scanning electron microscopy (SEM) of femoral condyle cartilage surface from control (Ctrl) and *cKO* mice. Images show comparable sGAG staining and collagen fibril network organization between genotypes following the reduction of decorin (*n* ≥ 3). d) AFM-nanoindentation shows comparable indentation modulus (*E*_ind_) between genotypes (*n* = 5). **e-f)** Comparison of e) Safranin-O histology and f) nanoindentation modulus (*E*_ind_) of condyle cartilage between male and female mice at 9 and 18 months (9M and 18M) for both genotypes (*n* ≥ 3 per genotype/sex). No significant sex-dependent differences were detected between male and female joints. Panels d and f: mean ± 95% CI, n.s., not significant. Each data point represents the average value of > 10 locations measured from one animal).

**Extended Data Figure 2.**
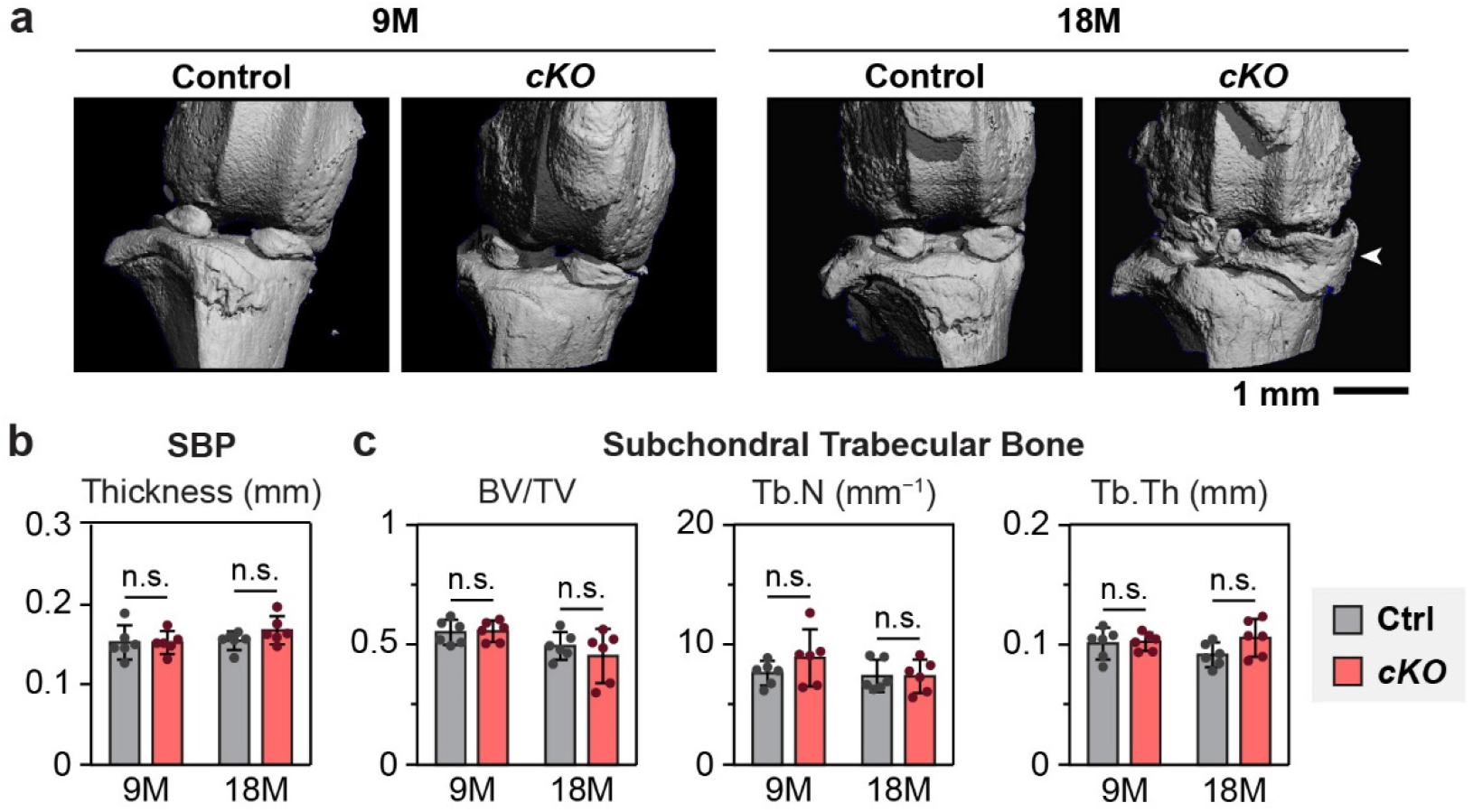
Micro-computed tomography (µCT) analysis of subchondral bone structure. **a)** Representative reconstructed 3D µCT images of knee joints from control (Ctrl) and *cKO* mice at 9 and 18 months (9M and 18M). Arrowhead indicates the increased calcification of meniscal anterior horn in *cKO* joint at 18M. **b-c)** Quantifications of b) subchondral bone plate (SBP) thickness, and c) subchondral trabecular bone structural parameters, including bone volume fraction (BV/TV), trabecular number (Tb.N), and trabecular thickness (Tb.Th). No significant differences were detected between genotypes at either age (mean ± 95% CI, n.s., not significant, *n* = 6). Each data point represents the average value measured from one animal).

**Extended Data Figure 3.**
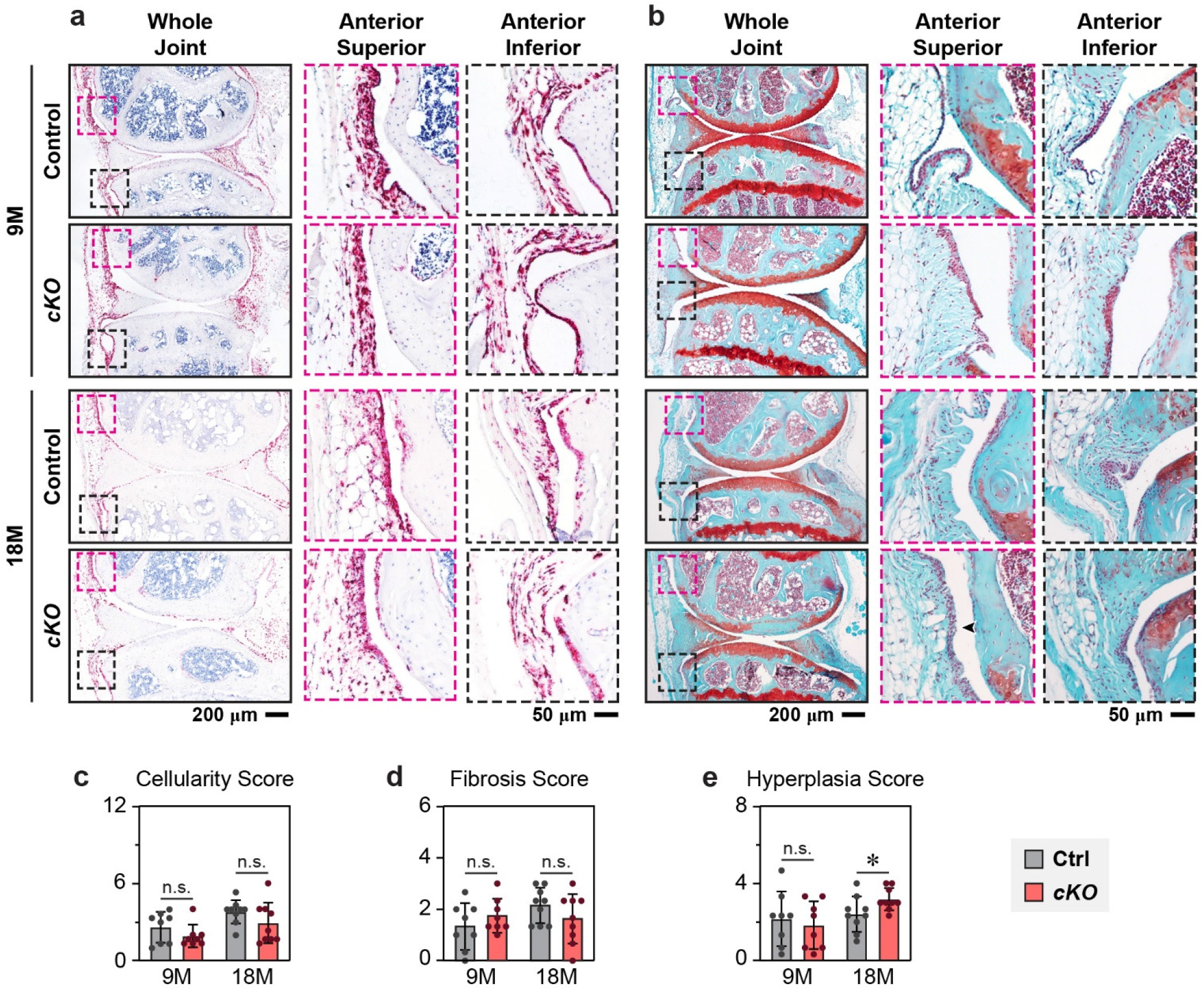
Decorin (*Dcn*) expression and structure of joint synovium. **a)** RNAscope *in situ* hybridization for *Dcn* and **b)** Safranin-O/Fast Green images of sagittal knee joint sections from control (Ctrl) and *cKO* mice at 9 and 18 months (9M and 18M). Whole-joint views and higher-magnification images of the anterior superior and anterior inferior synovium regions show comparable spatial distribution and intensity of *Dcn* expression between genotypes. Arrowheads indicate regions of synovial structural alterations. **c-e)** Histological scores of synovial morphology, including c) cellularity, d) fibrosis, and e) synovial lining hyperplasia (mean ± 95% CI, *: *p* < 0.05, n.s., not significant; *n* = 8). Each data point represents the average value measured from one animal.

**Extended Data Figure 4.**
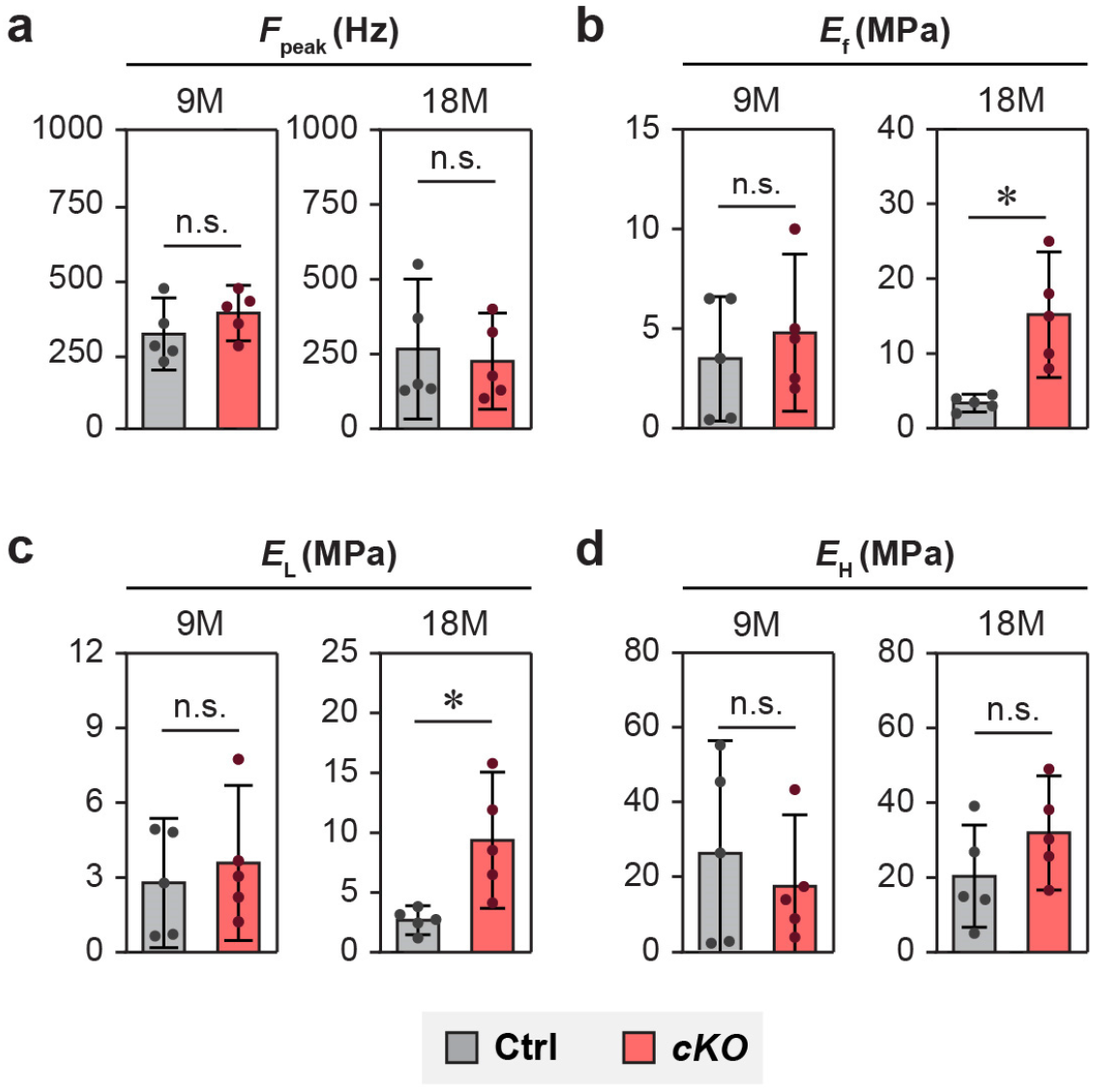
Additional poroelastic properties of *Dcn^cKO^* and control cartilage measured via the nanorheometric test. **a)** Peak frequency at the maximum phase angle (*F*_peak_), **b)** Tensile-only Young’s modulus (*E*_f_) derived from the fibril-reinforced poroelasticity model, **c,d)** Averaged dynamic modulus at the low frequency end (*E*_L_, ≤ 5 Hz), and d) the high frequency end (*E*_H_, 800-1,200 Hz) (mean ± 95% CI, *: *p* < 0.05, n.s.: not significant, *n* ≥ 5). Each data point represents the average value of > 10 locations measured from one animal.

**Extended Data Figure 5.**
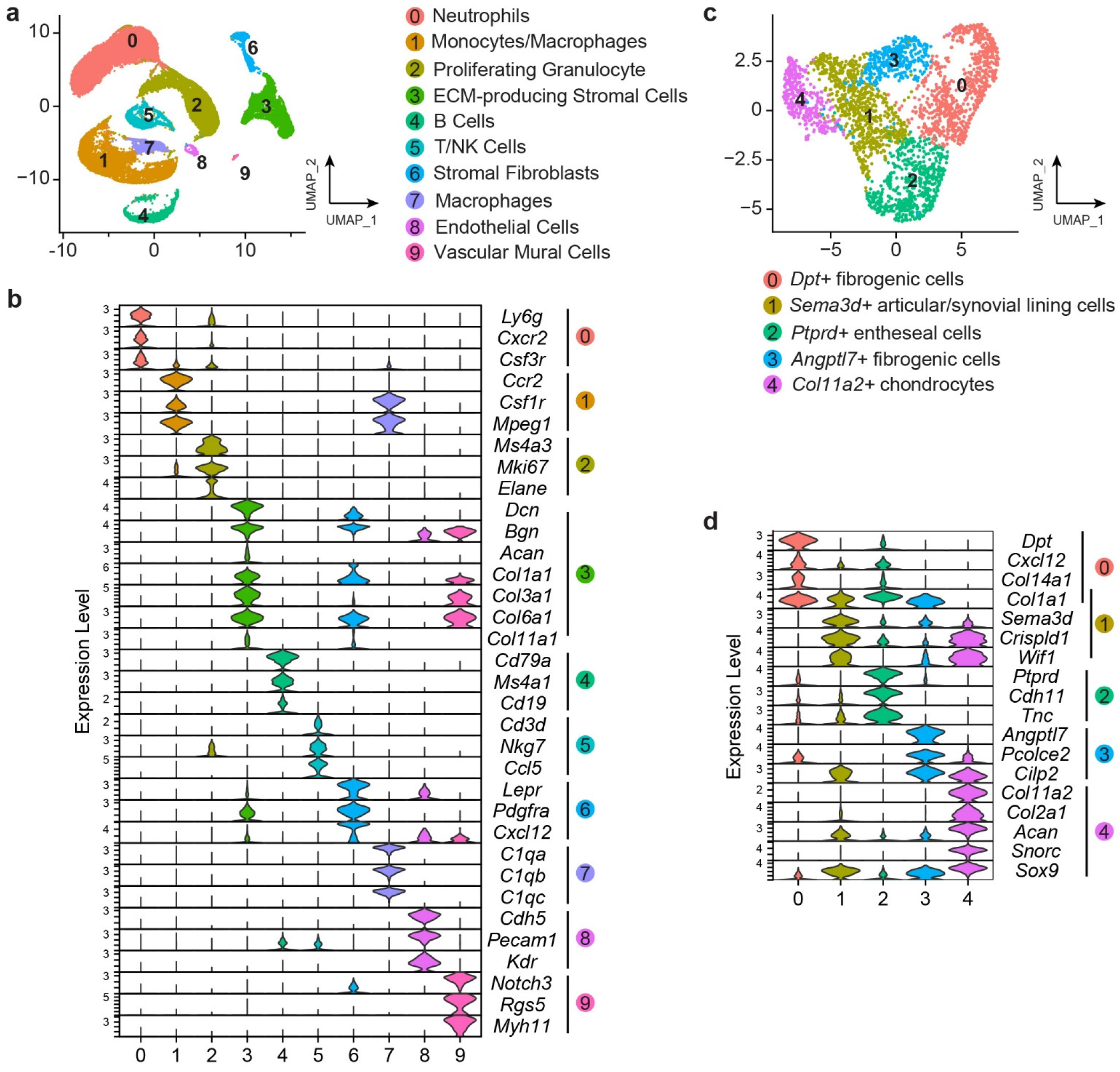
Cell clustering and sub-clustering analysis of scRNA-seq data at 9 months of age. **a)** UMAP projection of all cells isolated from 9-month-old control (Ctrl) and *cKO* tissues. Major cell populations are identified by unsupervised clustering and annotated based on cluster-enriched marker genes. Cluster 3 contains chondrocyte-associated ECM producing stromal cells and is selected for further analysis, as shown in Fig. 3. **b)** Violin plots showing expression of representative marker genes used for cell-type annotation across clusters. **c)** UMAP projection of cells from Cluster 3 following re-clustering and excluding irrelevant clusters, revealing transcriptionally distinct subpopulations. Subclusters are annotated based on DEGs and established lineage markers. **d)** Violin plots of sub-cluster-enriched marker genes validating subpopulation identities within the chondrocyte-associated compartment.

**Extended Data Figure 6.**
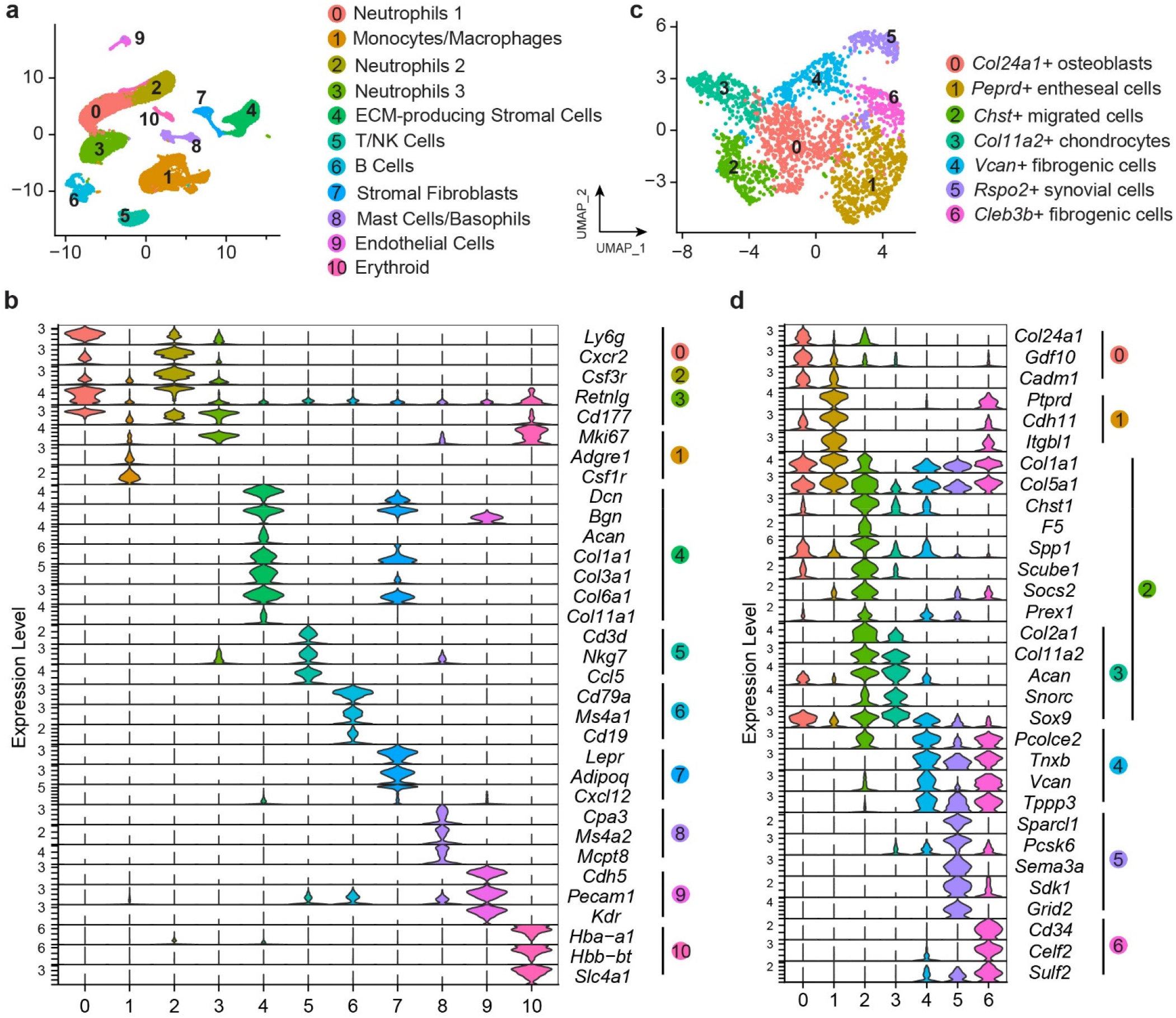
Cell clustering and sub-clustering analysis of scRNA-seq data at 18 months of age. **a)** UMAP projection of all cells isolated from 18-month-old control (Ctrl) and *cKO* tissues. Major cell populations are identified by unsupervised clustering and annotated based on cluster-enriched marker genes. Cluster 4 contains chondrocyte-associated ECM producing stromal cells and was selected for further analysis, as shown in Figs. 4 and 5. **b)** Violin plots showing expression of representative marker genes used for cell-type annotation across clusters. **c)** UMAP projection of cells from Cluster 4 following re-clustering and excluding irrelevant clusters, revealing transcriptionally distinct subpopulations. Subclusters are annotated based on DEGs and established lineage markers. **d)** Violin plots of subcluster-enriched marker genes validating subpopulation identities within the chondrocyte-associated compartment.

**Extended Data Figure 7.**
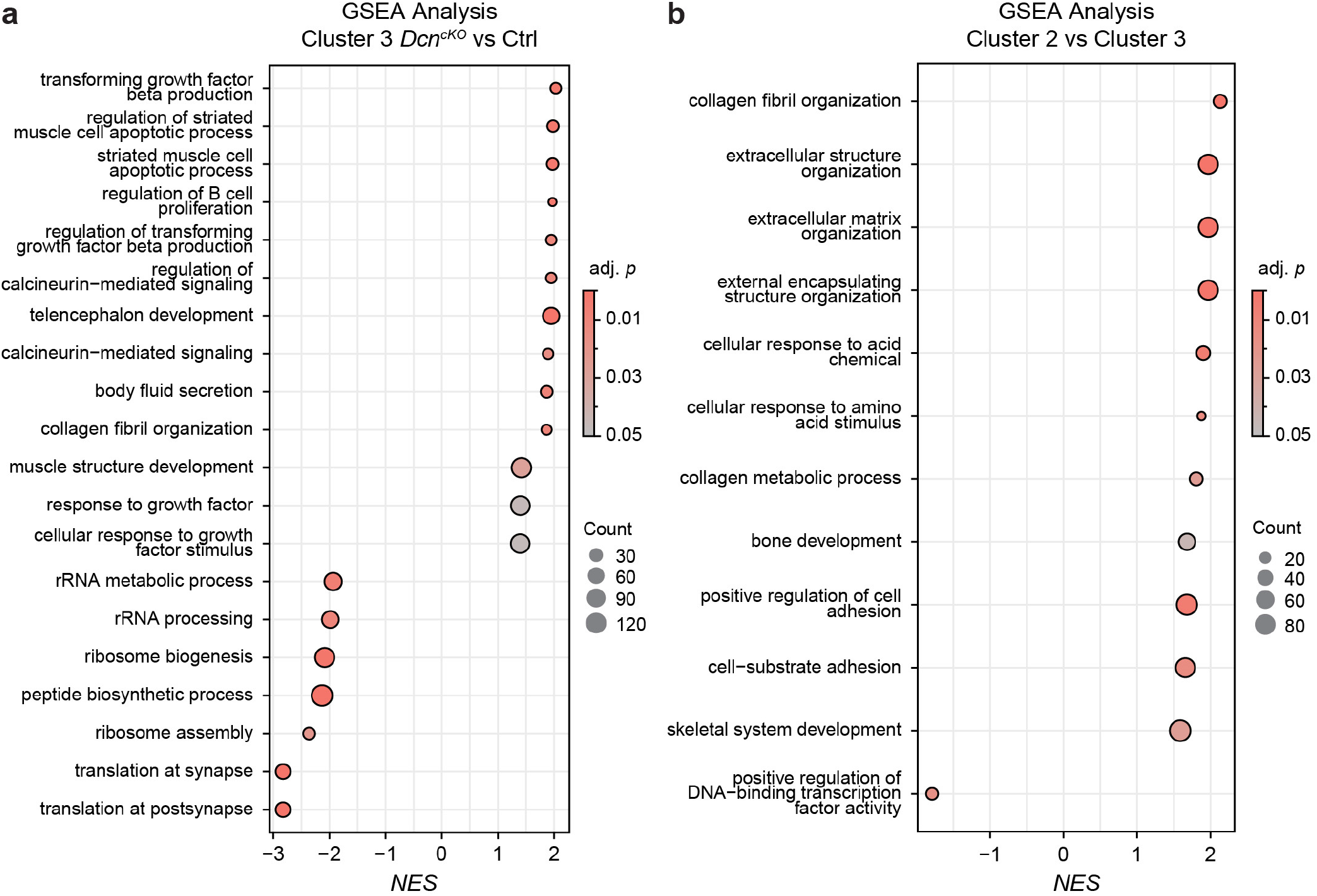
Gene set enrichment analysis (GSEA) of Cluster 3 chondrocyte and Cluster 2 emergent cell populations for scRNA-seq data at 18 months of age. **a)** GSEA of articular chondrocytes (Cluster 3) for *cKO* versus control (Ctrl) tissues. Top 20 significantly enriched biological processes (adjusted *p* < 0.05) are shown. **b)** GSEA of emergent cells (Cluster 2) versus articular chondrocytes (Cluster 3) for cells pooled from both genotypes. Significantly enriched processes (adjusted *p* < 0.05) highlight functional differences between the two populations. Processes are ranked by normalized enrichment score (NES). Dot size represents gene count, and color indicates adjusted *p*-value.

**Extended Data Figure 8.**
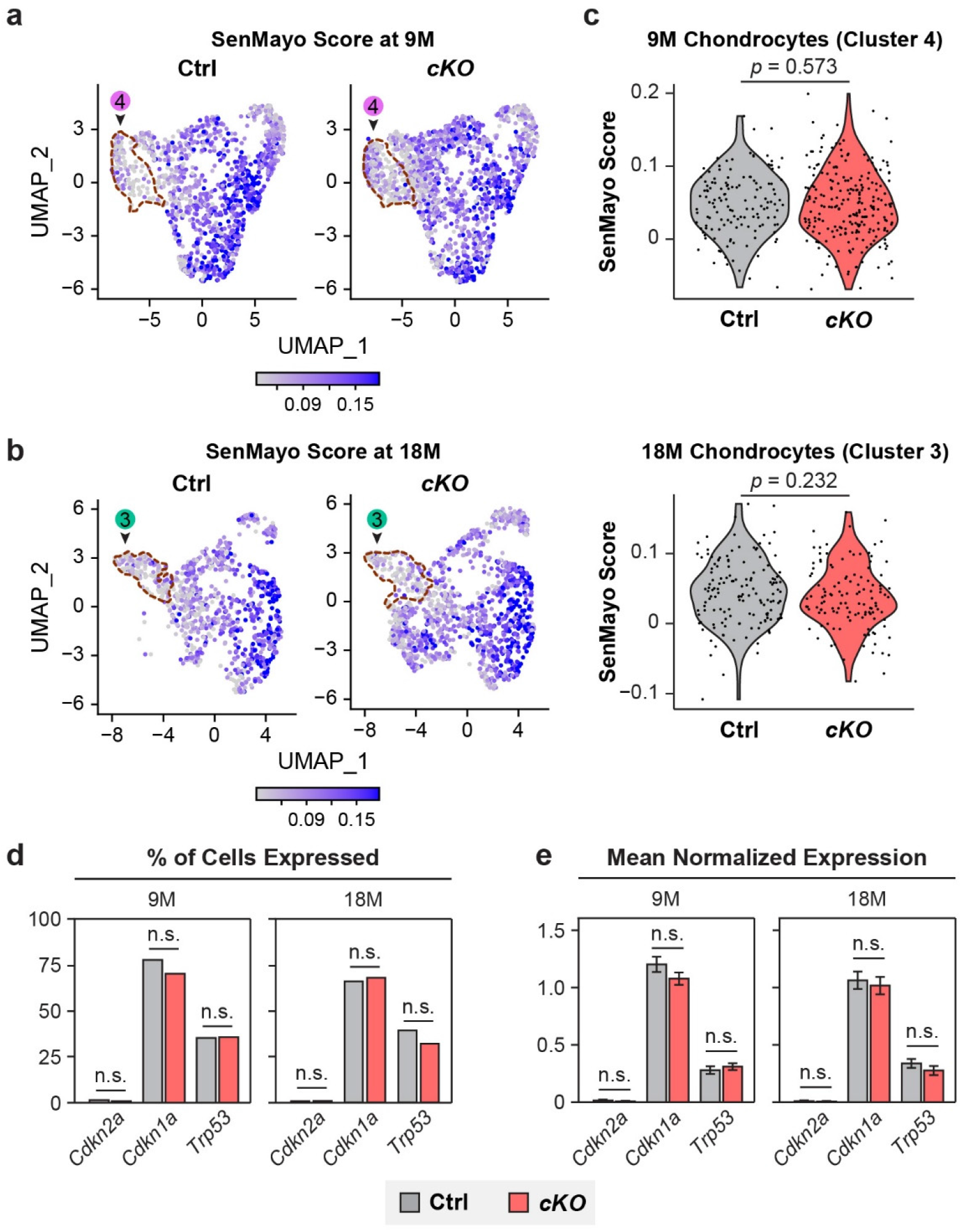
Articular chondrocytes exhibit similar senescence gene module scores between *Dcn^cKO^*and control cartilage. **a,b)** UMAP feature plots showing SenMayo module score distribution in a) 9-month-old and b) 18-month-old control (Ctrl) and *cKO* cartilage. The SenMayo gene signature comprises 115 senescence and SASP-associated genes, of which 108 and 109 were detected at 9M and 18M respectively. Articular chondrocyte clusters are highlighted by red outline and arrowhead (9M: Cluster 4 in Fig. 3a; 18M: Cluster 3 in Fig. 4a). **c)** Violin plots showing SenMayo module scores in articular chondrocytes at both ages. **d,e)** Comparisons of **d)** the percentage of articular chondrocytes expressing canonical senescence markers *Cdkn2a* (p16^INK4a^), *Cdkn1a* (p21^CIP1^) and *Trp53* (p53) and **e)** mean normalized expression of these markers (± SEM) between Ctrl and *cKO* chondrocytes. No significant differences were found between genotypes at either timepoint (*n* = 137 Ctrl and 229 *cKO* chondrocytes at 9M; *n* = 143 Ctrl and 123 *cKO* chondrocytes at 18M; n.s.: not significant).

**Extended Data Figure 9.**
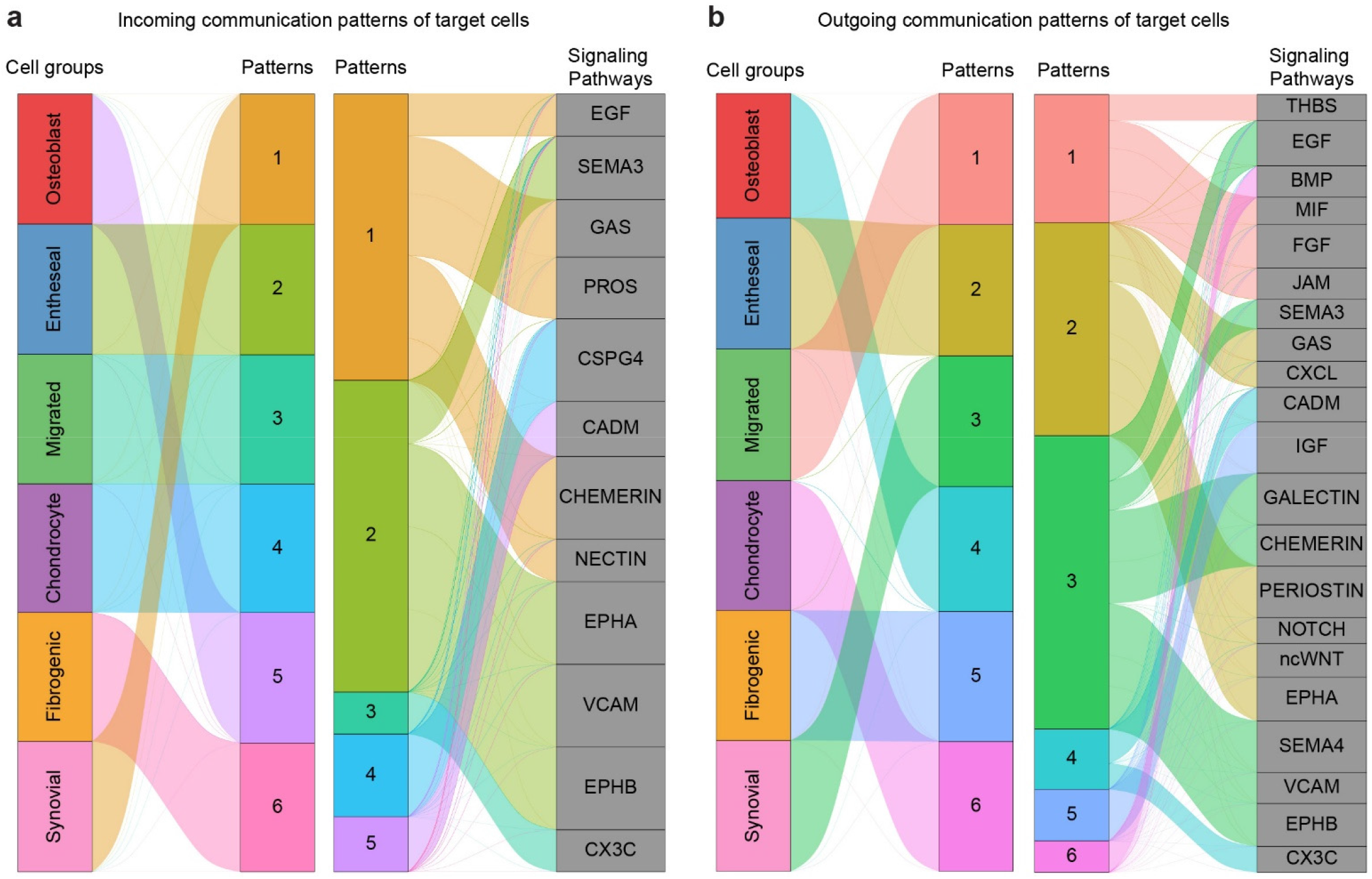
CellChat-inferred communication patterns and dominant signaling pathways for scRNA-seq data at 18 months of age. River (alluvial) plots showing **a)** incoming and **b)** outgoing communication patterns of target cells. Cell groups, as defined based on UMAP annotation in Fig. 4, are linked to communication patterns (middle columns) and their associated signaling pathways (right). Each stream is color-coded according to the pattern, and the stream thickness is proportional to aggregated communication probability by integrating contributions from ligand-receptor interactions. Patterns for which a given cell group contributes only minimally may appear visually absent from the pattern block in the second column due to insufficient weight.

**Extended Data Figure 10.**
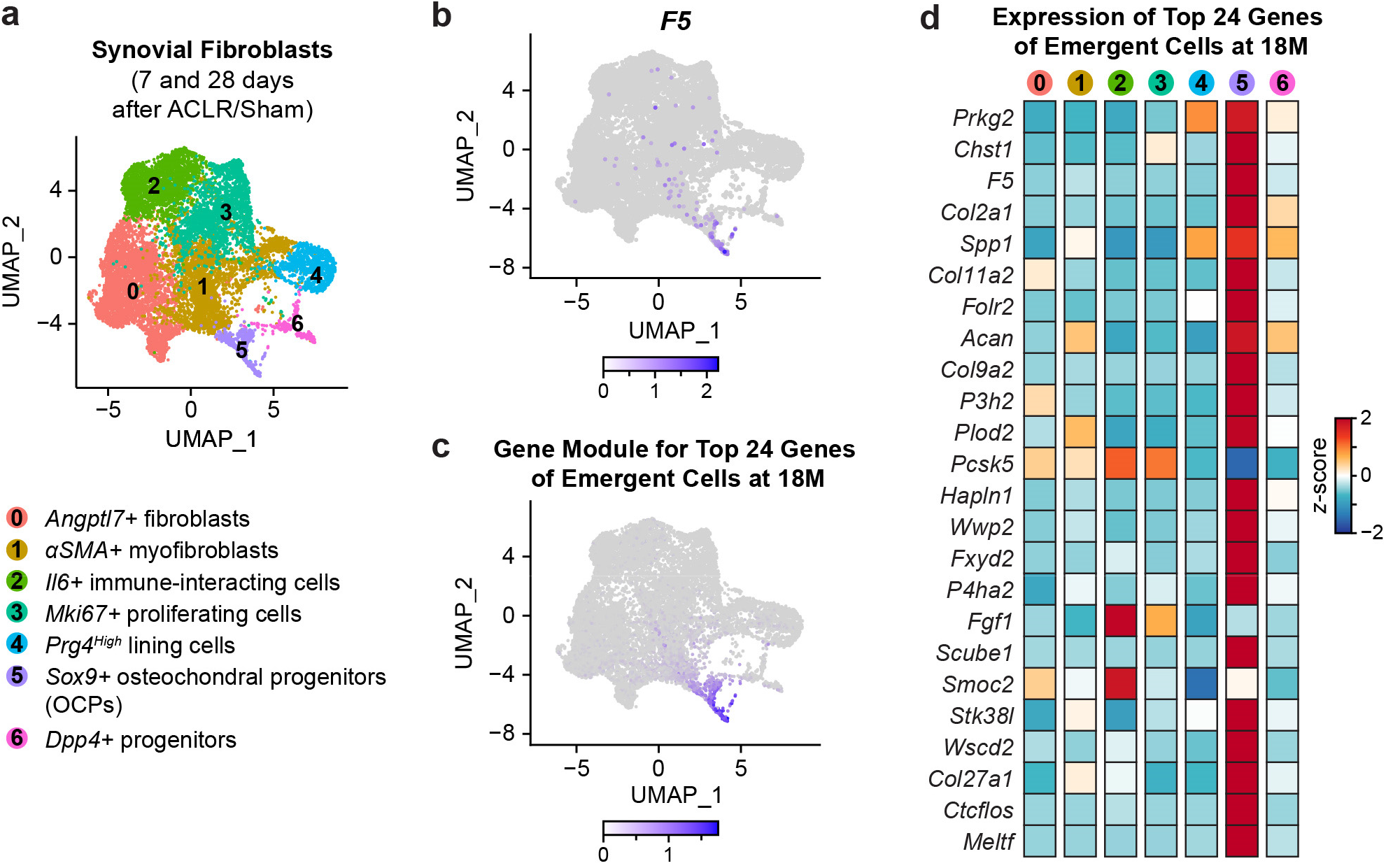
Emergent cell clusters in 18 months old *Dcn^cKO^* mice cartilage show gene markers implying osteochondral progenitors origin from synovium tissue. a) UMAP of previously described fibroblast subsets in murine synovium from Sham and anterior cruciate ligament rupture (ACLR)-operated joints (Data adapted from Ref. ^52^). Cluster 6 was identified as osteochondral progenitors. b) Feature plot of F5 expression. c) Gene module calculated from the top 24 marker genes identified by *FindAllMarkers* of the emergent cell cluster, described in Figure 5b. d) Heatmap shows the expression level of top 24 marker genes in synovium fibroblast subpopulations.

