## Supporting Materials for "Decorin-mediated Extracellular Matrix Stabilization Maintains Cartilage Integrity During Aging"

### SUPPLEMENTARY TABLES

**Table S1.** Summary of averaged values and statistical analysis outcomes of histological and μCT analysis on femoral condyle cartilage, subchondral bone and synovium for 9-month-old (9M) and 18-month-old (18M) decorin conditional knockout (*Dcn^cKO^*) and control (Ctrl) mice. Results are shown as mean ± 95% confidence interval (CI) from values averaged by each animal.

| | 9M | | | | | 18M | | | | | $p$-value (9M vs 18M) | |
|---|---|---|---|---|---|---|---|---|---|---|---|---|
| | Ctrl | | $Dcn^{cKO}$ | | $p$ | Ctrl | | $Dcn^{cKO}$ | | $p$ | Ctrl | $Dcn^{cKO}$ |
| | mean ± 95% CI | $n$ | mean ± 95% CI | $n$ | | mean ± 95% CI | $n$ | mean ± 95% CI | $n$ | | | |
| $t_{\mathrm{uncalcified}}$ (μm) | 32 ± 4 | 6 | 29 ± 7 | 6 | 0.451 | 32 ± 10 | 6 | 12 ± 10 | 6 | 0.008 | 0.822 | 0.010 |
| $t_{\mathrm{total}}$ (μm) | 105 ± 7 | 6 | 95 ± 10 | 6 | 0.079 | 101 ± 14 | 6 | 83 ± 13 | 6 | 0.072 | 0.527 | 0.172 |
| $\rho_{\mathrm{cell,\ uncalcified}}$ (× $10^{-3}$ cells/μm$^2$) | 5.8 ± 1.0 | 5 | 5.9 ± 0.5 | 5 | 1.000 | 5.2 ± 1.3 | 5 | 4.7 ± 3.0 | 5 | 1.000 | 0.626 | 0.626 |
| $\rho_{\mathrm{cell,\ calcified}}$ (× $10^{-3}$ cells/μm$^2$) | 2.4 ± 0.2 | 5 | 2.2 ± 0.3 | 5 | 0.300 | 1.8 ± 0.4 | 5 | 2.0 ± 1.2 | 5 | 0.728 | 0.008 | 0.630 |
| Mod. Mankin score | 0.06 ± 0.14 | 6 | 1.2 ± 1.2 | 6 | 0.061 | 0.8 ± 0.6 | 6 | 4.8 ± 1.1 | 6 | 0.004 | 0.015 | 0.004 |
| SBP Th (mm) | 0.15 ± 0.02 | 6 | 0.15 ± 0.01 | 6 | 0.982 | 0.16 ± 0.01 | 6 | 0.17 ± 0.02 | 6 | 0.360 | 0.485 | 0.232 |
| STB BV/TV | 0.56 ± 0.05 | 6 | 0.56 ± 0.05 | 6 | 0.870 | 0.50 ± 0.06 | 6 | 0.46 ± 0.11 | 6 | 0.842 | 0.108 | 0.108 |
| STB Tb.N (mm$^{-1}$) | 7.7 ± 1.0 | 6 | 9.0 ± 2.4 | 6 | 0.464 | 7.5 ± 1.4 | 6 | 7.4 ± 1.4 | 6 | 1.000 | 0.937 | 0.368 |
| STB Tb.Th (mm) | 0.10 ± 0.01 | 6 | 0.10 ± 0.01 | 6 | 0.811 | 0.09 ± 0.01 | 6 | 0.10 ± 0.02 | 6 | 0.166 | 0.358 | 0.641 |
| STB Tb.Sp (mm) | 0.12 ± 0.03 | 6 | 0.13 ± 0.04 | 6 | 0.877 | 0.16 ± 0.03 | 6 | 0.14 ± 0.03 | 6 | 0.798 | 0.080 | 0.395 |
| Syn. cellularity | 2.6 ± 1.0 | 8 | 1.9 ± 0.7 | 8 | 0.522 | 3.8 ± 0.7 | 9 | 2.9 ± 1.2 | 9 | 0.218 | 0.050 | 0.166 |
| Syn. fibrosis | 1.3 ± 0.8 | 8 | 1.8 ± 0.6 | 8 | 0.518 | 2.1 ± 0.5 | 9 | 1.6 ± 0.7 | 9 | 0.518 | 0.112 | 0.908 |
| Syn. hyperplasia | 2.2 ± 1.2 | 8 | 1.8 ± 1.0 | 8 | 0.831 | 2.4 ± 0.7 | 9 | 3.2 ± 0.4 | 9 | 0.084 | 0.583 | 0.068 |

**Table S2.** Summary of the distributions of collagen fibril diameter, $d_{\mathrm{col}}$, and von Mises concentration, $\kappa$, and statistical analysis outcomes of the femoral condyle cartilage surface for Ctrl and $Dcn^{cKO}$ mice at 9M and 18M, as measured by scanning electron microscopy (SEM) from $n = 4$ animals.

| | | 9M | | 18M | |
|---|---|---|---|---|---|
| | | Ctrl | $Dcn^{cKO}$ | Ctrl | $Dcn^{cKO}$ |
| $d_{\mathrm{col}}$ (nm) | mean | 45 | 66 | 44 | 51 |
| | 95% CI | [44 46] | [64 68] | [43 45] | [50 52] |
| | std | 15 | 22 | 12 | 15 |
| | $Q_1$ | 34 | 50 | 35 | 40 |
| | $Q_2$ | 42 | 64 | 44 | 49 |
| | $Q_3$ | 54 | 78 | 52 | 59 |
| | min | 19 | 20 | 16 | 27 |
| | max | 111 | 179 | 90 | 128 |
| $\kappa$ | mean | 0.11 | 1.88 | 0.19 | 1.22 |
| | 95% CI | [0.07 0.16] | [1.68 2.09] | [0.13 0.26] | [1.07 1.39] |
| | $n_{\mathrm{fibrils}}$ | 400 | 400 | 400 | 400 |

| | mean($d_{\mathrm{col}}$) | | var($d_{\mathrm{col}}$) | | $\kappa$ | |
|---|---|---|---|---|---|---|
| $p$-value (Ctrl vs $Dcn^{cKO}$) | 9M | 18M | 9M | 18M | 9M | 18M |
| | < 0.001 | < 0.001 | < 0.001 | 0.002 | < 0.001 | < 0.001 |
| $p$-value (9M vs 18M) | Ctrl | $Dcn^{cKO}$ | Ctrl | $Dcn^{cKO}$ | Ctrl | $Dcn^{cKO}$ |
| | 0.344 | < 0.001 | < 0.001 | < 0.001 | 0.686 | < 0.001 |

**Table S3.** Summary of averaged values and statistical analysis outcomes of the biomechanic properties of femoral condyle cartilage from *Dcn*$^{cKO}$ and *Ctrl* mice at 9M and 18M, shown as mean ± 95% CI from values averaged by each animal.

| | 9M | | | | | 18M | | | | | *p*-value (9M vs 18M) | |
|---|---|---|---|---|---|---|---|---|---|---|---|---|
| | Ctrl | | *Dcn*$^{cKO}$ | | *p* | Ctrl | | *Dcn*$^{cKO}$ | | *p* | | |
| | mean ± 95% CI | *n* | mean ± 95% CI | *n* | | mean ± 95% CI | *n* | mean ± 95% CI | *n* | | Ctrl | *Dcn*$^{cKO}$ |
| $E_{ind}$ (MPa) | 2.96 ± 1.04 | 9 | 4.22 ± 1.59 | 8 | 0.132 | 2.47 ± 0.95 | 12 | 11.86 ± 7.43 | 10 | 0.002 | 0.456 | 0.027 |
| $\delta_m$ (°) | 36 ± 13 | 5 | 32 ± 9 | 5 | 0.461 | 34 ± 7 | 5 | 21 ± 2 | 5 | 0.002 | 0.713 | 0.030 |
| $F_{peak}$ (Hz) | 317 ± 121 | 5 | 387 ± 92 | 5 | 0.476 | 267 ± 234 | 5 | 227 ± 161 | 5 | 0.704 | 0.609 | 0.043 |
| $E_H/E_L$ | 7.0 ± 4.3 | 5 | 4.4 ± 1.5 | 5 | 0.156 | 7.5 ± 2.6 | 5 | 3.7 ± 0.9 | 5 | 0.026 | 0.778 | 0.522 |
| $k$ (×$10^{-15}$ $m^4/(N·s)$) | 0.46 ± 0.30 | 5 | 0.85 ± 0.21 | 5 | 0.018 | 0.66 ± 0.42 | 5 | 4.3 ± 1.5 | 5 | 0.004 | 0.316 | 0.006 |
| $E_m$ (MPa) | 1.36 ± 1.30 | 5 | 1.71 ± 1.35 | 5 | 0.615 | 1.36 ± 0.59 | 5 | 4.40 ± 2.58 | 5 | 0.029 | 1.000 | 0.034 |
| $E_f$ (MPa) | 3.49 ± 3.12 | 5 | 4.8 ± 3.94 | 5 | 0.490 | 3.40 ± 1.19 | 5 | 15.20 ± 8.39 | 5 | 0.034 | 0.943 | 0.028 |
| $E_L$ (MPa) | 2.78 ± 2.60 | 5 | 3.58 ± 3.11 | 5 | 0.599 | 2.70 ± 1.21 | 5 | 9.39 ± 5.69 | 5 | 0.029 | 0.943 | 0.038 |
| $E_H$ (MPa) | 26.4 ± 30.1 | 5 | 17.4 ± 19.1 | 5 | 0.505 | 20.4 ± 13.6 | 5 | 32.0 ± 15.3 | 5 | 0.310 | 0.628 | 0.274 |

**Table S4.** Summary of the distributions of collagen fibril length, $l_{col}$, fibril diameter, $d_{col}$, and von Mises concentration, $\kappa$, and statistical analysis outcomes for collagen II fibrils deposited on glass substrate through microfluidics channel with or without the addition of decorin (collagen II:decorin at 25:1 and 15:1, w/w), as imaged by SEM and quantified through CT-FIRE. Data are collected from > 4,500 fibrils pooled from three independent experiments for each condition.

| | | Collagen II only | 25:1 | 15:1 |
|---|---|---|---|---|
| $l_{col}$ (μm) | mean | 1.29 | 0.94 | 0.94 |
| | 95% CI | [1.27 1.32] | [0.92 0.95] | [0.92 0.96] |
| | std | 0.86 | 0.63 | 0.61 |
| | $Q_1$ | 0.73 | 0.54 | 0.55 |
| | $Q_2$ | 1.03 | 0.74 | 0.74 |
| | $Q_3$ | 1.58 | 1.10 | 1.09 |
| | min | 0.39 | 0.33 | 1.09 |
| | max | 8.67 | 6.39 | 7.25 |
| $d_{col}$ (nm) | mean | 71 | 61 | 57 |
| | 95% CI | [70.4 71.1] | [60.8 61.7] | [56.2 56.9] |
| | std | 12 | 16 | 11 |
| | $Q_1$ | 63 | 52 | 49 |
| | $Q_2$ | 71 | 58 | 56 |
| | $Q_3$ | 78 | 66 | 63 |
| | min | 35 | 32 | 29 |
| | max | 148 | 217 | 110 |
| $\kappa$ | mean | 2.68 | 0.85 | 0.98 |
| | 95% CI | [2.60 2.76] | [0.81 0.89] | [0.94 1.02] |
| | $n_{fibrils}$ | 4500 | 4500 | 4500 |

| | $l_{col}$ | | | $d_{col}$ | | | $\kappa$ | | |
|---|---|---|---|---|---|---|---|---|---|
| | Col II only vs 25:1 | Col II only vs 15:1 | 15:1 vs 25:1 | Col II only vs 25:1 | Col II only vs 15:1 | 15:1 vs 25:1 | Col II only vs 25:1 | Col II only vs 15:1 | 15:1 vs 25:1 |
| *p*-value | < 0.001 | < 0.001 | 0.975 | < 0.001 | < 0.001 | < 0.001 | < 0.001 | < 0.001 | 0.003 |

**Table S5.** Summary of the distributions of collagen fiber length, $l_{col}$, and von Mises concentration, $\kappa$, and statistical analysis outcomes for collagen II fibers formed between pairs of chondrocyte spheroids with or without the addition of decorin (collagen II:decorin at 25:1 and 15:1, w/w), as imaged by polarizing light microscope (PLM) and quantified through CT-FIRE. Outcomes for $\kappa$ were also analyzed based on both pooled values and averaged values from individual spheroid pairs. Data are collected from ≥ 950 fibers pooled from ≥ 5 spheroid pairs across three independent experiments.

| | | By individual fibers | | |
|---|---|---|---|---|
| | | Collagen II only | 25:1 | 15:1 |
| $l_{col}$ (μm) | mean | 69 | 58.9 | 56.1 |
| | 95% CI | [66.5 72.0] | [57.2 60.7] | [55.1 57.0] |
| | std | 43.3 | 32.9 | 27.7 |
| | $Q_1$ | 38.8 | 37.1 | 36.9 |
| | $Q_2$ | 54.7 | 47.7 | 47.4 |
| | $Q_3$ | 84.3 | 70.5 | 65.8 |
| | min | 30.0 | 30.0 | 30.0 |
| | max | 293.9 | 278.7 | 282.5 |
| $\kappa$ | mean | 2.37 | 1.36 | 1.11 |
| | 95% CI | [2.21 2.53] | [1.27 1.45] | [1.06 1.16] |
| | $n_{fibers}$ | 943 | 1317 | 3311 |
| $\kappa$ (by spheroid pairs) | Mean | 2.93 | 1.59 | 1.27 |
| | 95% CI | [2.03 3.83] | [0.42 2.77] | [0.87 1.66] |
| | $N_{spheroid\ pairs}$ | 7 | 5 | 8 |

| | $l_{col}$ | | | $\kappa$ | | | $\kappa$ (by spheroid pairs) | | |
|---|---|---|---|---|---|---|---|---|---|
| | Col II only vs 25:1 | Col II only vs 15:1 | 15:1 vs 25:1 | Col II only vs 25:1 | Col II only vs 15:1 | 15:1 vs 25:1 | Col II only vs 25:1 | Col II only vs 15:1 | 15:1 vs 25:1 |
| $p$-value | < 0.001 | < 0.001 | 0.016 | < 0.001 | < 0.001 | < 0.001 | 0.028 | 0.002 | 0.754 |

**Table S6.** Summary of averaged values and statistical analysis outcomes of the indentation modulus, $E_{ind}$, measured on bovine cartilage surface with and without the infiltration of decorin. Values were obtained from 12 explant plugs from 4 animals per group with 10-15 indentation locations per half-plug.

| $E_{ind}$ (MPa) (mean ± 95% CI) | Decorin Infiltration | | $p$-value (Decorin + vs −) |
|---|---|---|---|
| | ( − ) | ( + ) | |
| Untreated | 0.091 ± 0.045 | 0.134 ± 0.075 | 0.012 |
| sGAG-depleted | 0.024 ± 0.006 | 0.058 ± 0.026 | 0.002 |
| $p$-value | 0.012 | 0.017 | |

**Table S7.** Information for single-cell RNA sequencing datasets. Raw data (FASTQ files) and filtered barcodes, features, and matrix files are deposited in GEO under the file names listed below. Samples 1 to 4 were collected and sequenced together following identical sample preparation protocols, with each sample pooled from one male and one female mouse prior to cell loading by the sequencing core; these four samples were merged for downstream analysis. Samples 5 to 7 correspond to 18-month timepoint femur tissue. Sample 5 and 7 were collected alongside Samples 1 to 4. Sample 6 was collected from one female mouse in a separate pilot sequencing run, and Sample 7 was collected from one male mouse to balance sex representation for the 18-month $Dcn^{cKO}$ group. Spearman correlation analysis confirmed high similarity in cell composition between Samples 6 and 7 within the cartilage-related matrix-producing cluster (R = 0.97), and Samples 5 to 7 were therefore merged for downstream analysis. For the 18-month timepoint, only femur samples were collected and analyzed to avoid masking the observed site-specific phenotypic differences between femur and tibia (Figure 1, 2). For the 9-month timepoint, both femur and tibia samples were collected and pooled to more comprehensively capture the transcriptomic effects of decorin conditional knockout on cartilage cell function.

| Sample number | GEO number | File name | Description in manuscript |
|---|---|---|---|
| 1 | | LinHan-20240111-GEX-9M-femur-control | 9-month control |
| 2 | | LinHan-20240111-GEX-9M-tibia-control | 9-month control |
| 3 | | LinHan-20240111-GEX-9M-femur-Cko | 9-month $Dcn^{cKO}$ |
| 4 | | LinHan-20240111-GEX-9M-tibia-Cko | 9-month $Dcn^{cKO}$ |
| 5 | | LinHan-20240412-GEX-18M-femur-control | 18-month control |
| 6 | | LinHan-20230518-GEX-18M-femur-Cko | 18-month $Dcn^{cKO}$ |
| 7 | | LinHan-20240412-GEX-18M-femur-Cko-2ed | 18-month $Dcn^{cKO}$ |

### SUPPLEMENTARY METHOHDS

#### Detailed Methods for single-cell RNA Sequencing Data Processing and Analysis

**Data Loading and Quality Control.** Filtered feature-barcode matrices generated by *Cell Ranger* were imported into *R (v4.5.2)* using *Read10X()* and converted into *Seurat objects (v5.4.0)* using *CreateSeuratObject()*, excluding genes expressed in fewer than 10 cells and cells with fewer than 100 detected features. Mitochondrial gene content was quantified per cell using *PercentageFeatureSet()* with the pattern "^mt-". Quality control was performed on each sample independently. The lower nCount_RNA cutoff was determined empirically by computing the proportion of cells removed across thresholds ranging from 500 to 10,000 (increments of 500) and identifying the trough following the first peak of the numerical derivative curve, yielding sample-specific cutoffs of 2,500 for 9-month femur and 2,250 for 9-month tibia and all 18-month samples. Cells were retained if they satisfied all the following criteria: 200 < nFeature_RNA < 93rd percentile of the sample distribution; nCount_RNA > sample-specific lower cutoff and < 97th percentile; and mitochondrial gene content < 5%. Upper bounds were defined dynamically per sample to account for sequencing depth variation and remove likely multiplets.

**Normalization and Merging.** Each sample was independently normalized using *NormalizeData()* (log normalization, scale factor = 10,000), and the top 3,000 highly variable features were identified using *FindVariableFeatures()*. QC-filtered Seurat objects were merged using *merge()* with unique cell ID prefixes to preserve sample-of-origin information, and RNA layers were consolidated into a single unified layer using *JoinLayers()*.

**Cell Cycle Scoring, Scaling and Dimensionality Reduction.** Cell cycle phase was assigned to each cell using *CellCycleScoring()* based on curated S-phase and G2M-phase gene lists, generating S and G2M scores. Data were scaled using *ScaleData()*, regressing out S score, G2M score, and mitochondrial ratio to reduce confounding variation from cell cycle state and mitochondrial gene expression. Highly variable features were reidentified using *FindVariableFeatures()*, and PCA was performed using *RunPCA()* on the resulting variable features.

**Cell Annotation and Subpopulation Selection.** Following PCA, UMAP and unsupervised clustering were performed using dims = 1:10 and resolution = 0.1 for the 9-month dataset. Cluster marker genes were identified using *FindAllMarkers()* (adjusted $p$-value ≤ 0.05). To assist annotation, pct.1 and pct.2 were calculated to prioritize genes highly expressed within a cluster but absent in others, filtered across four specificity thresholds (60/40, 70/30, 80/20). Cell type identity was assigned based on this specificity-filtered marker list in combination with canonical marker gene expression, and immune and blood cell contaminants were removed. Cluster 4 was identified as the matrix-producing population based on expressions of *Dcn*, *Bgn*, *Acan*, *Col1a1*, *Col3a1*, *Col6a1*, and *Col11a1* (Extended Data Figure 6) and isolated using *subset()* for sub-clustering. Sub-clustering was first performed using dims = 1:25 and resolution = 0.2, which revealed Cluster 5 to be transcriptionally distinct and spatially isolated from all other cells, distorting the UMAP embedding; this cluster was therefore excluded. Sub-clustering was repeated on the remaining cells using dims = 1:10 and resolution = 0.3, and the resulting clusters constituted the cartilage-related matrix-producing population carried forward for all downstream analyses (Fig. 3 and Extended Data Figure 6).

For the 18-month dataset, UMAP and unsupervised clustering were performed using dims = 1:25 and resolution = 0.1. Cluster marker genes were identified and annotated using the same approach described above. Cluster 4 was again identified as the matrix-producing population based on expressions of *Dcn*, *Bgn*, *Acan*, *Col1a1*, *Col3a1*, *Col6a1*, and *Col11a1* (Extended Data Figure 7) and isolated for sub-clustering. Sub-clustering was first performed using dims = 1:10 and resolution = 0.2, which revealed Cluster 7 to be transcriptionally distinct and spatially isolated, distorting the UMAP embedding; this cluster was therefore excluded. Sub-clustering was repeated using dims = 1:10 and resolution = 0.3, and the resulting clusters constituted the matrix-producing population carried forward for downstream analyses (Fig. 4 and Extended Data Figure 7).

**Differential Expression and Visualization.** Differentially expressed genes (DEGs) between *cKO* and control conditions were identified per cluster using *FindMarkers()*, with *cKO* set as ident.1 and control as ident.2, such that positive avg_log2FC values reflect upregulation in *cKO* relative to control. DEGs were considered significant at adjusted *p*-value < 0.05. Cluster marker expression was visualized using violin plots (*VlnPlot()*), dot plots (*DotPlot()*), and feature plots *(FeaturePlot()*), and DEGs were visualized using volcano plots with significantly upregulated and downregulated genes highlighted by adjusted *p*-value and log2 fold-change thresholds. All plots were generated using Seurat's built-in functions and *ggplot2*.

**Gene Set Enrichment Analysis.** GSEA was performed using *gseGO()* from the *clusterProfiler* package (v4.18.3), with Gene Ontology Biological Process (GO:BP) terms annotated against the mouse genome database (*org.Mm.eg.db, v3.22.0*). Genes were pre-ranked using a signed metric combining direction and significance: rank = sign(log2FC) × (−log10(*p*-value)), ordering genes from most significantly upregulated to most significantly downregulated. The following parameters were applied: pvalueCutoff = 0.05, minGSSize = 15, maxGSSize = 500, nPermSimple = 10000, and eps = 1e-300. A random seed was set prior to analysis for reproducibility (*set.seed(1)*). Semantic similarity between GO terms was assessed using *godata()* from the *GOSemSim package (v2.36.0)*. Results were ranked by absolute normalized enrichment score (|NES|), and pathways relevant to extracellular matrix organization, collagen, and TGF-β signaling were selected for visualization as dot plots using *dotplot*.

**Cell-Cell Communication Analysis.** Intercellular communication was inferred using *CellChat (v1.6.1).* A CellChat object was constructed using *createCellChat()* from the raw count matrix and cell metadata, with cells grouped by cluster identity. The mouse ligand-receptor interaction database *(CellChatDB.mouse)* was applied, and overexpressed signaling genes and ligand-receptor interactions were identified using *identifyOverExpressedGenes()* and *identifyOverExpressedInteractions().* Mouse protein-protein interaction data were incorporated using *projectData().* Communication probabilities were computed using *computeCommunProb()* with the triMean method, and interactions involving clusters with fewer than 10 cells were removed using *filterCommunication()*. Pathway-level communication scores were aggregated using *computeCommunProbPathway()* at a significance threshold of 0.05. Dominant outgoing and incoming communication patterns were identified by non-negative matrix factorization using *identifyCommunicationPatterns(),* with the number of patterns determined separately for each direction using *selectK(),* selecting the value at which both Cophenetic and average silhouette scores began to decline, yielding 6 outgoing and 6 incoming patterns. Relationships between cell populations, communication patterns, and signaling pathways were visualized as alluvial plots using *netAnalysis_river()* from the *ggalluvial package (v0.12.5).*

**SenMayo Module Score Analysis.** The SenMayo mouse gene set was obtained from published dataset (Saul et al., *Nat. Commun.* 2022, 13:4287), comprising 118 senescence and SASP-associated genes. Gene coverage was assessed prior to module score computation and genes absent from the dataset expression matrix were excluded from scoring. Coverage exceeded 91% at both timepoints, above the recommended threshold for reliable module scoring. Module scores were computed using *AddModuleScore()* in Seurat (v5.4.0) with seed = 42, applied to the RNA assay following *JoinLayers()*. The average expression of the detected SenMayo genes is relative to a set of randomly sampled control genes, where positive scores indicate above-average senescence gene expression and negative scores indicate below-average expression relative to the transcriptome background. Normalized expression values for *Cdkn2a*, *Cdkn1a*, and *Trp53* were extracted from articular chondrocyte subsets using *GetAssayData()*. For each gene, the percentage of cells with expression > 0 (percent expressed) and mean normalized expression with standard error of the mean (SEM) calculated per condition. Comparisons were performed using the Mann-Whitney U test for mean expression and Fisher's exact test for proportions of cell expressing the markers.
